# SorCS2 binds progranulin and regulates motor axon outgrowth

**DOI:** 10.1101/2023.02.08.527602

**Authors:** PB Thomasen, A Salašová, H Login, S Beel, J Tranberg-Jensen, P Qvist, PL Ovesen, S Nolte, LN Nejsum, MV Chao, J Dasen, P Van Damme, K Kjaer-Sorensen, C Oxvig, A Nykjaer

## Abstract

Motor neuron development requires an orchestrated action of trophic factors and guidance cues for axons to reach their targets. Here, we identify SorCS2 as a novel receptor for progranulin (PGRN) that is required for motor axon outgrowth in zebrafish and mice. In both species motor neurons express SorCS2, and PGRN is produced in cells juxta-positioned the projecting axon, but in mice the neurons also co-express PGRN. In zebrafish, *sorcs2* knockdown produces stunted and aberrantly branched motor axons, and in *Sorcs2^-/-^* mice, forelimb innervation and motor neuron regeneration are substantially perturbed; phenotypes also observed in fish and mice lacking PGRN. SorCS2 binds PGRN and while motor neuron cultures from wildtype mice respond to exogenous PGRN by axon outgrowth, knockout neurons are unresponsive. Remarkably, when co-expressed in the same cells, SorCS2 controls secretion of PGRN. We conclude that SorCS2 navigates motor neuron development and enables axon regeneration through binding of PGRN.

## Introduction

Motor neurons are the final relay that transmits signals from the central nervous system to skeletal muscles in the periphery, thereby enabling muscle contraction and movement. Their development is complex and requires a concerted action of trophic factors and guidance cues that secures neuronal survival, axonal outgrowth and the proper formation of functional neuromuscular junctions (NMJs). This process relies on signaling molecules expressed by the motor neurons themselves as well the surrounding cells in a manner that is highly controlled in time and space^1,2^. The same neurotrophic factors and guidance cues are often reactivated during axonal regeneration after peripheral nerve injury, as they support neuronal survival and navigate the axonal regrowth following Wallerian degeneration^3^. Though motor axon outgrowth has been extensively studied in both mice and zebrafish, the proteins and pathways involved are still not fully understood.

The Vps10p-domain (Vps10p-d) receptor, SorCS2, is a member of the sortilin receptor family comprising sortilin, SorLA, SorCS1, and SorCS3. The receptors predominate in distinct neuronal populations of the central- and peripheral nervous system where they bind a vast number of ligands and engage in cell signaling, endocytosis, and intracellular trafficking^4,5^. The expression of SorCS1, −2, and −3 dynamically changes during embryonic development, suggesting important roles in defining neuronal structures^6–11^. Conceptually, signaling through a Vps10p-d receptor requires formation of heteromeric complexes with the ligand and a co-receptor^5,12,13^. By doing so, SorCS2 has been implicated in activities ranging from growth cone retraction, to regulation of neuronal plasticity and apoptosis of peripheral glia^13–17^.

All Vps10p-d receptors are initially produced as a proform containing a propeptide that is cleaved off in the late Trans-Golgi Network (TGN) to generate a mature single-chain receptor. However, SorCS2 stands out from the other family members as it can be further converted from the single-chain form that controls growth cone retraction and synaptic plasticity, into a two-chain variant that is proapoptotic when expressed in glia cells^13,16^. The sorting activities of the receptor family are many and each receptor may engage in several trafficking pathways to control ligand secretion, endocytosis, recycling, and lysosomal sorting from the TGN. Notably, SorCS2 can establish the surface exposure and synaptic targeting of membrane receptors such as the tyrosine receptor kinase, TrkB, and NMDA receptors^16–18^.

PGRN is a multifunctional protein produced by several cell types including microglia and neurons^19^. Among other functions, it promotes neuronal development and repair, sustains neuronal integrity, and suppresses neuroinflammatory responses^20–22^. For instance, motor neuron development is substantially perturbed in zebrafish lacking PGRN and in *Grn* knockout mice regeneration of injured motor neurons is delayed^23–27^. The neuroprotective effect of PGRN is further supported by *GRN* haploinsufficiency being a common cause of frontotemporal dementia^19,28,29^. Molecularly, PGRN exerts its pleiotropic functions by two mechanisms. It can signal cross the plasma membrane, and it can sustain lysosomal function by providing mature granulins and by targeting the enzyme prosaposin to lysosomes^20,21,27,30^. Whereas the pathways that mediate lysosomal sorting have been extensively studied^31–33^, the receptor(s) responsible for binding PGRN and transducing its neurotrophic activity remains elusive.

Here we show that SorCS2 regulates motor neuron outgrowth in fish and mice. In both species, SorCS2 inactivation resulted in stunted motor neurons, and in SorCS2 knockout (*Sorcs2^-/-^*) mice regeneration of injured adult motor neurons was perturbed too. Further, we demonstrate that SorCS2 binds PGRN to facilitate its cellular signaling and secretion. Our findings identify SorCS2 as a novel PGRN receptor and highlight the importance of this interaction for motor neuron development and regeneration.

## Results

### *Sorcs2* is expressed in motor neurons in the ventral spinal cord of zebrafish embryos

To interrogate the function of SorCS2 in motor neuron development we first assessed its embryonic expression in zebrafish. *sorcs2* was weakly expressed in embryos from 12 hours post fertilization (hpf) and increased during development up to 48 hpf (**Fig. 1A**). We subsequently analysed its tissue expression by alkaline *in situ* hybridisation (ISH) at 24 hpf, a time point when many mechanisms regulating motor neuron development are active (**Fig. 1B**). *sorcs2* was detected in the ventral spinal cord corresponding to the location of motor neurons as well as in muscle cells within the somites (**Fig. 1B**). To enhance resolution of the ISH we employed third generation *in situ* hybridization chain reaction (*in situ* HCR v3.0)^34^ on transgenic fish expressing GFP driven by the motor neuron-specific promotor *mnx1 (Tg(mnx1:GFP)^ml2^)*. In accordance with the regular ISH, we detected *sorcs2* mRNA in motor neuron cell bodies in the spinal cord and in muscle in the trunk at 24 hpf as shown by lateral view (**Fig. 1C-D**) and in cross sections (**Fig. 1E-F**). Analysis for *sorcs2* expression using a single-cell transcriptome atlas of zebrafish embryos^35^ (**Fig. S1A-C**) confirmed our *in situ* HCR data.

**Figure 1:**
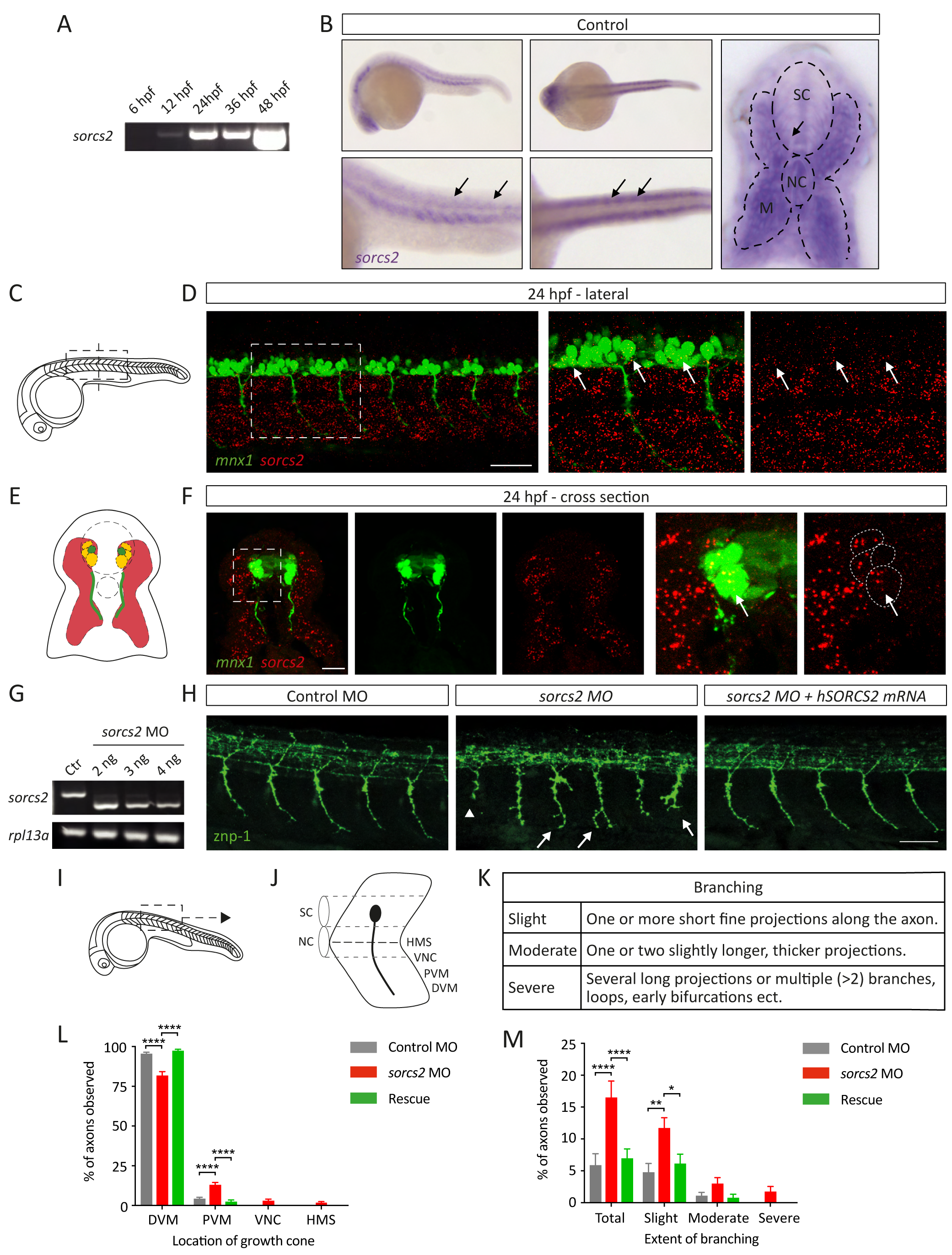
*Sorcs2* knockdown causes stunted and aberrantly branched primary motor axons in zebrafish embryos. **(A)** RT-PCR showing s*orcs2* is increasingly expressed from 12 hours post-fertilization (hpf). **(B)** I*n situ* hybridization (ISH) of *sorcs2* at 24 hpf. Left; Lateral and dorsal view of control embryos showing staining in somites and ventral spinal cord (SC) Right: cross section of control SC. Muscle (M) outlined with dashed lines. NC = notochord. Motor neurons are indicated by arrows. **(C-F)** *In situ* HCR for *sorcs2* in *Tg(mnx1:GFP)^ml2^* embryos at 24 hpf showing *sorcs2* in GFP+ motor neurons (arrows) in lateral (C-D) and cross sectional (E-F) view. (**C**) Area studied in **(D**) is boxed. (**E**) Representation of cross sections shown in (**F**). Images are z-projection of 4 μm (D) and 10 μm (F). Marked areas in D and F, left panels, are enlarged in the corresponding right panels, scalebar = 50 μm for (D) and 20 μm for (F). **(G)** Efficacy of splice-site inhibiting *sorcs2* morpholino (MO) injected at different dosages compared to 5 ng of control (ctr) MO assessed 24 hpf by RT-PCR of *sorcs2* and the housekeeping gene, *rpl13a.* Injections made with 5 ng p53 MO. **(H)** Znp-1 immunostaining showing abnormal length (arrowhead) and branching (arrows) of CaP axons in zebrafish treated with sMO compared to control MO at 27 hpf. Rescue of phenotype by co-injection of sMO with h*SORCS2* mRNA. Scalebar = 50 μm. **(I)** Representation of zebrafish embryo. The box shows area used for CaP analysis. **(J)** Model showing CaP projection into four defined regions used for scoring axon outgrowth^71^. HMS, horizontal myoseptum; VNC, ventral edge of notochord; PVM and DVM, distal and proximal ventral myotome. **(K)** Definitions for scoring branching^72^. Axon outgrowth **(L)** and branching **(M)** in CaP from control MO (n = 21) and *sorcs2* MO zebrafish (n = 27), and following rescue (n = 18) are shown in percentage. Error bars: s.e.m., n = number of embryos. P-value determined with two-way ANOVA and Tukey’s multiple comparisons. **** < 0.0001, ** < 0.01.

### *Sorcs2* knockdown causes stunted and aberrantly branched motor axons

To study the impact of SorCS2 deficiency on motor neuron outgrowth, we knocked down (KD) receptor expression by injecting a morpholino oligomer that disrupts a splice site in sorcs2 (sMO). At a dosage of 3.7 ng, the sMO effectively perturbed sorcs2 splicing at 24 hpf (Fig. 1G) with no effect on developmental rate or morphology (Fig. S2). We assessed motor neuron length and branching in sMO-injected embryos at 27 hpf by whole-mount immunofluorescence (IF) staining using a znp-1 antibody that labels Synaptotagmin 2 (Fig. 1H). Zebrafish have three primary motor neurons innervating each somatic hemisegment, where the axon of the caudal primary motor neuron (CaP) is the first to grow, extending its projection into the ventral somite as indicated in Fig. 1I-J. In control-injected embryos, 95.6% of the CaP axons reached the distal ventral myotome (DVM), and all axons extended beyond the notochord (Fig. L). In contrast, axons of sorcs2 KD embryos were stunted with 81.9% of axons reaching the DVM and 1.9% only reaching the horizontal myoseptum (HMS) (p < 0.0001). In addition, 16.5% of the axons were branched in sorcs2 KD larvae compared to 5.9% in control (p < 0.0001), and severe branching was only observed in the sMO-injected group (Fig. 1M). Confirming the causative effect of sorcs2 knockdown on these phenotypes, co-injection with the human SORCS2 mRNA fully rescued the phenotype to the extent that 97.5% of the axons now exhibited normal growth (p < 0.0001) and 7% displayed branching (p < 0.0001) (Fig. 1L-M). We further analysed motor neuron length in sorcs2 knockout embryos (sorcs2-/-) from heterozygous cross of the sorcs2sa11642 fish line that harbors a disruptive mutation in exon 10. Similar to KD larvae, sorcs2-/- embryos exhibited stunted CaP axons as only 83.4% of axons reached the DVM compared to 97% in wildtype (wt) siblings (p = 0.022) (Fig. S3A). Injection of the sMO into the sorcs2-/- embryos did not further aggravate the motor neuron phenotype (Fig. S3B), which validates that the sMO is specific and has no off-target effects. Because of poor breeding of the sorcs2sa11642 fish, we continued our studies using the KD model.

To determine the dynamics of the outgrowth of the CaP axons in *sorcs2* deficient embryos, we subjected *Tg(mnx1:GFP)^ml2^* zebrafish to time-lapse imaging. CaP axons exit the spinal cord around 16-18 hpf and reach the HMS around 20-24 hpf where they make the first synaptic contacts. They pause here for a few hours before continuing ventrally into the myotome^36^. In *sorcs2* KD embryos axons paused for a longer period at the HMS though they retained their ability to extend beyond the choice point and grow ventrally (**Fig. 2A, Suppl. Movie 1-2**). However, several axons remained truncated and some started branching prematurely at later stages.

### Perturbed acetylcholine receptor clustering and motility in *sorcs2* KD embryos

Given the aberrant motor axon outgrowth in *sorcs2* deficient embryos, we asked whether axons were still capable of forming functional synapses with the acetylcholine receptors (AChRs) at the NMJ. To address this, co-staining of the motor axons and AChR clusters was performed at 27 and 48 hpf. During normal development, a prepattern of AChR clusters are partly replaced by neuromuscular synapses at 27 hpf. AChRs are clustered beneath outgrowing motor axons but are not yet precisely colocalized with clusters of synaptic vesicles in nascent presynaptic terminals.

Further, diffuse non-clustered AChRs are present at the yet non-innervated myoseptum, which separates adjacent myotomes^37^. This pattern we observed in control-injected embryos in which motor axons co-localized with AChR clusters (**Fig. 2B**). In marked contrast, AChR clusters in the *sorcs2* KD embryos were elongated and diffuse thereby resembling the pattern of immature, non-innervated clusters present at earlier stages of development. Similarly, co-localization between axons and AChRs were incomplete (arrows, **Fig. 2B**). Co-injection with human *SORCS2* mRNA fully rescued the phenotype. At 48 hpf motor axons have normally extended their branches into laterally located muscle fibres and turned at the edge of the myotome into the myosepta^37^. We found that in *sorcs2* KD embryos, axons had advanced less into the myosepta both in the ventral and dorsal direction. Furthermore, the co-localization between axon branches and AChRs in the somites remained reduced (asterisks, **Fig. 2B**). At this stage, *in situ* HCR on *Tg(mnx1:GFP)^ml2^* embryos showed that *sorcs2* was still highly expressed in motor neurons (**Fig. 2C-D**).

**Figure 2:**
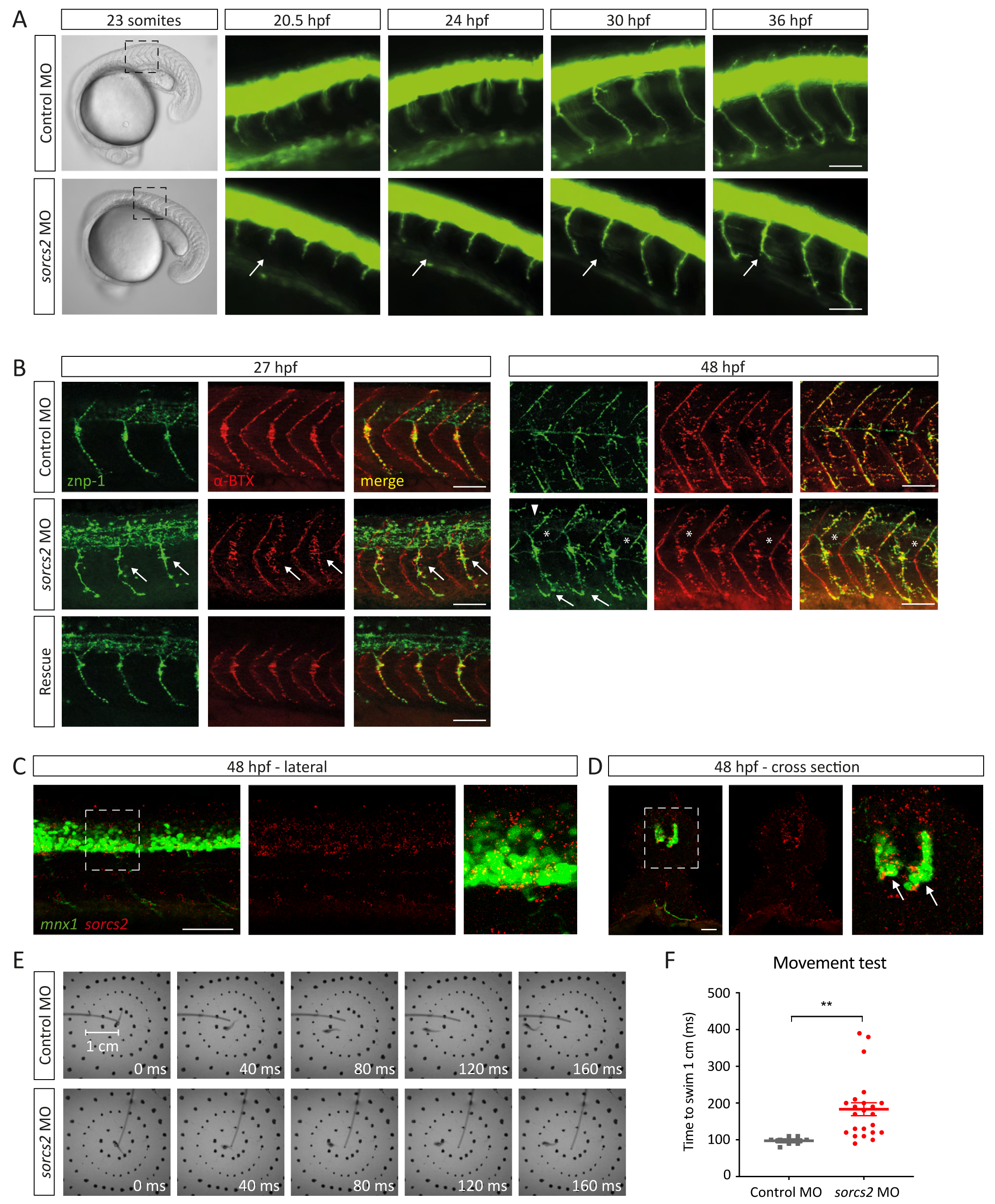
Neuromuscular defects and altered motility in *sorcs2* deficient embryos. **(A)** Differential interference contrast images of 20.5 hpf embryos and time-lapse of control and *sorcs2* MO *Tg(mnx1:GFP)^ml2^* from 20.5-36 hpf (enlargement of black box). The white arrow shows a truncated axon, which pauses longer at the first choice point (HMS) before continuing ventrally, yet remaining truncated at later stages. Scalebar = 20μm. **(B)** Immunofluorescent staining of the NMJ. Lateral view of znp-1 (axons, green) and bungarotoxin (AChRs, red) in control MO, *sorcs2* MO and rescue embryos at 27 hpf (left) and 48 hpf (right). At 27 hpf, AChR clusters are elongated, diffuse and without complete co-localization with axons in *sorcs2* MO (arrows). At 48 hpf, axons in *sorcs2* KD embryos have advanced less into the myosepta both at the ventral (arrows) and dorsal sides (arrow head, no innervation) and with less co-localization between axon branches and AChRs in somite (asterisks). Scalebar = 50 μm. *In situ* HCR of 48 hpf *Tg(mnx1:GFP)^ml2^* embryos showing *sorcs2* in GFP+ motor neuron cell bodies with lateral view **(C)** and cross section **(D)**. Images are z-projection of 16 μm (C) and 10 μm (D). Dashed squares are enlarged on the right side in each panel, scalebar = 50 μm for (C) and 20 μm for (D). **(E)** Images show responses of 48 hpf control and *sorcs2* MO embryos to touch. The distance between the center and second circle is 1 cm. **(F)** Time spent to swim 1 cm. *Sorcs2* MO embryos shows a significantly slower escape response to touch. Error bars: s.e.m. within groups. P-value determined with student’s t-test, ** < 0.01.

Prompted by the impaired synapse development, we next assessed the embryonic motility. We used a touch response assay^38^ in which the motility of 48 hpf embryos is assayed as the time it takes to swim 1 cm after a light touch. Here, the *sorcs2* KD embryos showed a significantly slower escape response and they spent on average 183 ms to swim 1 cm compared to the control embryos that only used 98 ms (**Fig. 2E-F**, p = 0.008, **Suppl. Movie 3-4**). Taken together our results demonstrated that SorCS2 is required for proper motor neuron development, NMJ maturation, and motor function in zebrafish.

#### Impaired motor axon outgrowth in *Sorcs2^-/-^* mice

Given the phenotype in zebrafish and that SorCS2 is highly conserved among species, we speculated that SorCS2 might also be critical for motor neuron development in mice. Murine, spinal motor neurons develop from progenitor cells located in the medial neural tube. They progress into postmitotic neurons around E9.5 and migrate laterally to position themselves into specific motor columns according to the muscle fibres they innervate^1,39^. Motor axons exit the spinal cord mainly through the ventral roots and innervate their target muscle around E13-E14^1,40^. Accordingly, we first determined the expression pattern of SorCS2 in E12.5 mouse embryos by IF staining (**Fig. 3A-C**). At the brachial level, SorCS2 was highly expressed in the floor plate and the ventral spinal cord where motor neurons are located, whereas staining was absent in the dorsal spinal cord and dorsal root ganglion that contain sensory neurons. At this age SorCS2 was also present in skeletal muscle, oesophagus, trachea, and boundary cap cells lining the entry and exit points of the peripheral nerve roots (**Fig. 3A**). Co-staining for Hb9 and Foxp1, markers that specify most of the spinal motor neurons and designate the limb-innervating lateral column of motor neurons at brachial and lumbar levels^1,41^, respectively, showed prominent co-expression with SorCS2 (**Fig. 3A-C**).

**Figure 3:**
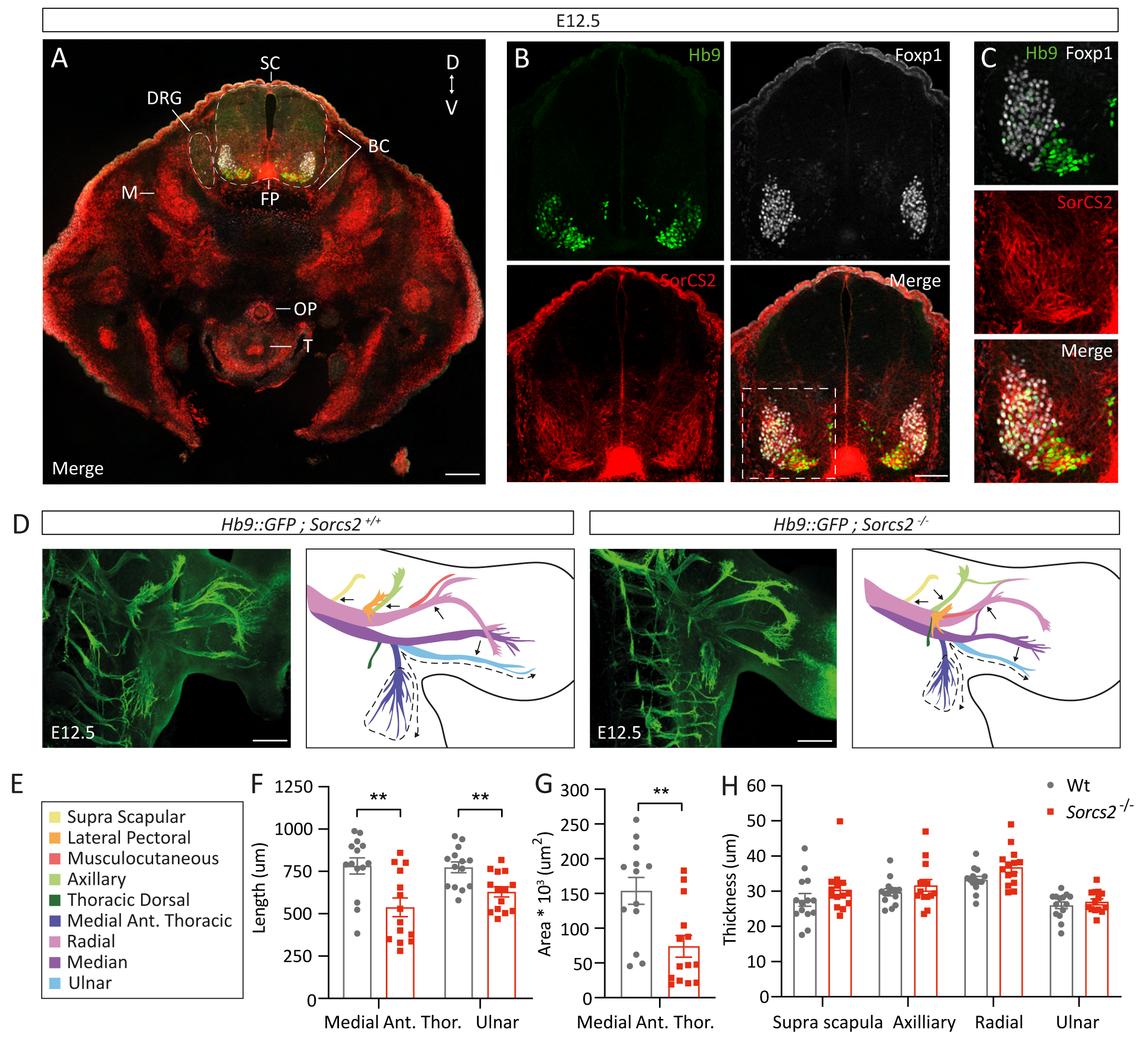
Perturbed motor axon outgrowth in *Sorcs2^-/-^* mice. **(A-C)** Immunofluorescent staining of SorCS2 (red) and motor neuronal transcription factors, Hb9 (green) and Foxp1 (white) in horizontal sections of E12.5 mice. SorCS2 is expressed in motor neurons, floor plate (FP), muscle (M), oesophagus (OE), trachea (T) and boundary cap (BC) cells, but not, or only very faintly, in dorsal root ganglia (DRG). Heart, lungs and intestines have been removed. D = dorsal, V = ventral. Scalebar = 200 μm in (A) and 50 μm in (B). **(C)** Enlargement of the area of interest boxed in (B) shows expression of SorCS2 in motor neurons of the ventral horn. **(D)** GFP fluorescence of whole-mount *Hb9::GFP* and *Sorcs2^-/-^*;*Hb9::GFP* embryos at E12.5. Axon outgrowth is impaired in SorCS2-deficient embryos. Schematic representations of the innervation pattern with motor nerves color coded according to Catela *et al.* 2016^74^. Dashed lines indicate the area measured, and dashed arrows and arrows the nerves of which the length and diameter have been quantified, respectively. Scalebar = 200 μm. **(E)** Color-codes of the forelimb nerves. Quantification of nerve lengths **(F)**, the area covered by the medial anterior thoracic nerve **(G),** and the nerve diameters **(H)** for control and *Sorcs2^-/-^* embryos (n =14). Data shown as mean ± s.e.m. P-value determined with multiple t-test, Holm-Šídák corrected, or Mann-Whitney test for area spanned, ** < 0.01.

Given the expression of SorCS2 in Hb9- and Foxp1-postive motor neurons, we asked whether *Sorcs ^-^* mice, like the zebrafish, display innervation deficits. Hence, knockout mice were crossed with *Hb9::GFP* transgenic mice, and innervation of the forelimbs and trunk was visualized by whole-mount imaging for GFP (**Fig. 3D-E**). Notably, in the absence of SorCS2, axonal outgrowth was substantially compromised. The ulnar nerve was 18% shorter in the SorCS2-deficient animals (**Fig. 3F**, wt: 774 μm, *Sorcs2^-/-^*: 630 μm, p = 0.0048), and the medial anterior thoracic nerve that innervates the cutaneous maximus and anterior latissimus dorsi muscles was 31% shorter compared to wt mice (wt: 783 μm, *Sorcs2^-/-^*: 539 μm, p = 0.0048). Further, the total area covered by the terminal axon arbors of the medial anterior thoracic nerve was reduced by no less than 52% (wt: 153,763 μm^2^, *Sorcs2^-/-^*: 73,984 μm^2^, p = 0.0034) (**Fig. 3G**). In contrast, SorCS2 expression did not impact the thickness of the motor nerves (**Fig. 3H**). We conclude that SorCS2 is required for proper motor neuron development also in mice.

### Binding of progranulin to SorCS2 and their expression in zebrafish and mice

We tested a panel of proteins implicated in axon pathfinding and trophic support for binding to SorCS2 using surface plasmon resonance (SPR) analysis and cellular binding assays. We failed to detect any consistent binding of EphrinA1-5, EphrinB1-3, EphA1-8, EphB1-6, UNC5H4, and netrin-1 in either of the assays (data not shown). However, PGRN bound robustly to the extracellular domain of SorCS2 by SPR analysis with an affinity of ∼3 nM (**Fig. 4A**). This was intriguing since knockdown of the PGRN-A encoding gene (*grna*) in zebrafish phenocopies the truncation defects and premature branching of motor neurons present in the *sorcs2* KD embryos^23–25^. Hence, we speculated that SorCS2 could be required to control PGRN function. To test this hypothesis, we first probed PGRN expression during motor neuron development in zebrafish and mice. Though the expression was generally sparse, *in situ* HCR revealed *grna* mRNA in zebrafish embryos in cells along the yolk extension most likely corresponding to immune cells (arrow heads, **Fig. 4B-C**). Strikingly, mRNA transcripts were also present in cells located in close proximity to the motor neurons (arrows, **Fig. 4B-C**). At 24 hpf, *grna* transcripts were most abundant juxtapositioned the initial segment of the projecting motor axons (**Fig. 4B**), whereas at 27 hpf *grna* was now located more distally lining the motor axons in both the ventral and dorsal myotome (**Fig. 4C-E**). Analysis of deposited single-cell transcriptome data^35^ suggested that the *grna* expressing cells were accounted for by macrophages and neutrophils. In accordance with our *in situ* HCR data, *Grna* was not detectable in motor neurons (**Fig. S4A-C**).

**Figure 4:**
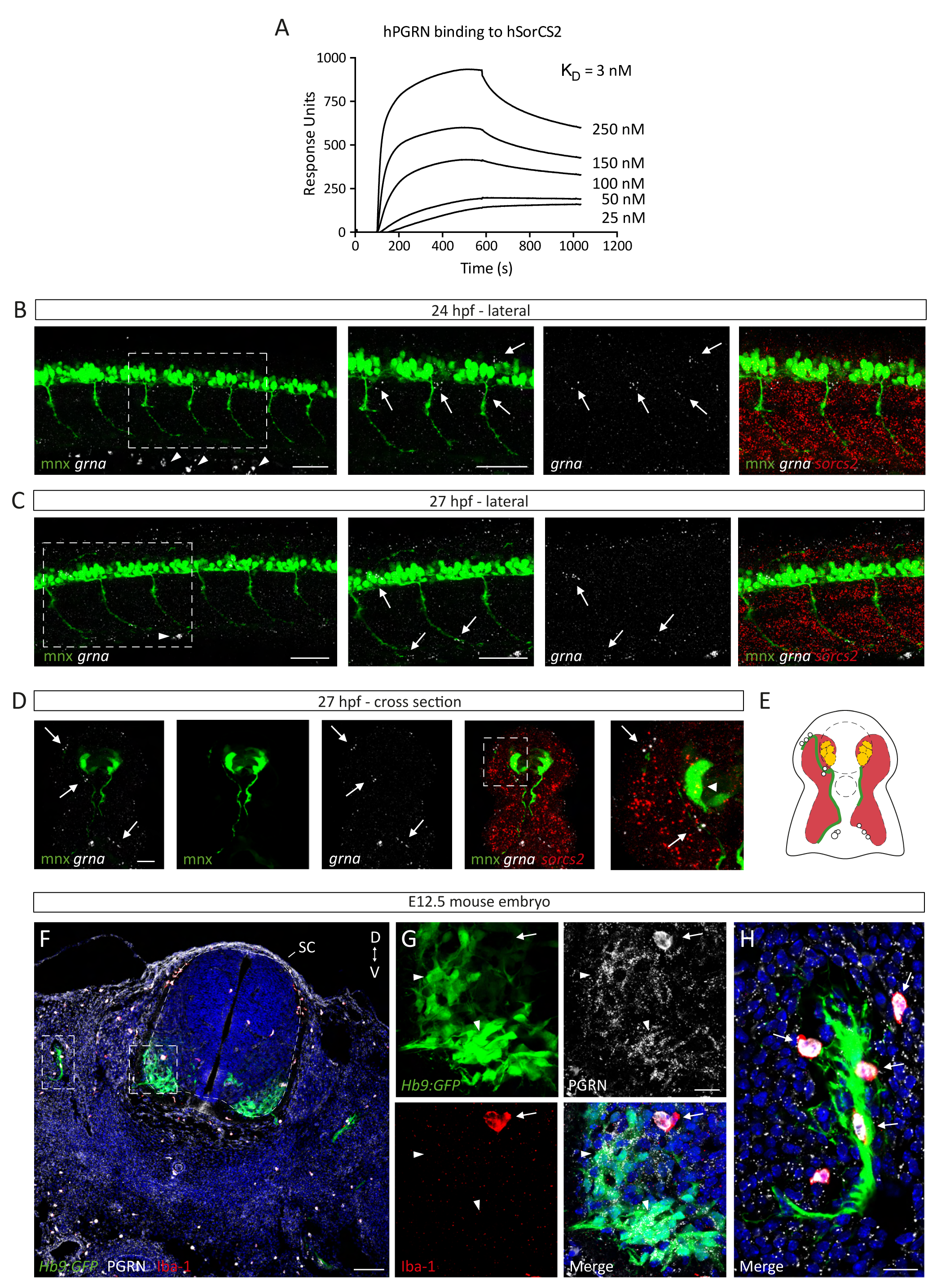
Binding of progranulin to SorCS2 and expression in zebrafish and mice. **(A)** Concentration-dependent binding of hPGRN to immobilized SorCS2 (Kd = 3 nM) demonstrated by surface plasmon resonance analysis. *In situ* HCR on *Tg(mnx1:GFP)^ml2^* embryos for *grna* shows expression along the yolk extension likely representing immune cells (arrow heads) and in close proximity to motor axons (arrows) at 24 **(B)** and 27 hpf **(C-D)**. Images are z-projection of 2 μm (B), 14 μm (C), and 10 μm (D). Dashed squares are enlarged to the right, scalebar = 50 μm for (B-C) and 20 μm for (D). **(E)** Schematic representation of a cross section of 27 hpf depicting *sorcs2* expression (red) in motor neuron cell bodies and skeletal muscle and *grna* (white) along extending motor axons (green). **(F-H)** Immunofluorescent staining of PGRN (white) and macrophage/microglia-marker, Iba-1 (red), in horizontal sections of E12.5 *Hb9::GFP* mice. **(F)** PGRN is expressed in scattered, brightly stained cells of which many co-express Iba-1 throughout the embryo. SC = spinal cord, V = ventral, D = dorsal. **(G)** Enlargement of ventral horn (white square) showing PGRN in Iba-1+ cells closely positioned to motor neurons (arrow), and PGRN expression in developing GFP+ motor neurons (arrow head). **(H)** Enlargement of projecting motor nerve (white rectangle) with PGRN in Iba-1+ cells (arrows) in close proximity to the motor axons (green). Nuclei are stained using Hoechst. Scalebar = 100 μm in (F) and 20 μm in (G-H).

We next studied PGRN expression in sections of the spinal cord from E12.5 *Hb9::GFP* mice by IF microscopy (**Fig. 4F-H**). We found that PGRN, like SorCS2 (cf. **Fig. 3A-C**), was present in Hb9-postive neurons (arrow heads, **Fig. 4G**). RNA scope ISH confirmed this expression pattern and further demonstrated that *Sorcs2* transcripts are abundant in the motor neurons too (**Fig. S5**).

Interestingly, some cells scattered throughout the embryo showed particularly bright PGRN labelling. IF microscopy revealed these cells to co-express Iba-1, which designates microglia and macrophages (**Fig. 4F-H**). Strikingly, outside the spinal cord PGRN expression was enriched in Iba-1-positive macrophages flanking the projecting motor axon (arrows, **Fig. 4H**). Taken together the data suggests that progranulin inside the motor neurons or secreted by cells surrounding the growing axon may enable motor development through binding to SorCS2.

### PGRN-induced motor neurite outgrowth requires SorCS2 expression

We next compared neurite outgrowth and sprouting in primary motor neurons isolated from E13.5^42^ wt and *Sorcs2^-/-^* mice cultured for four days in the absence or presence of exogenous PGRN (1.5 nM) or BDNF (0.4 nM) as a positive control. Labelling for *β*-tubulin was used to visualize neurites and quantify their length and branching (**Fig. 5A**). Whereas we did not observe any effect on the number of branches in wt neurons following PGRN stimulation (**Fig. 5B**), total neurite length was increased by 46% from 156 μm to 227 μm (p = 0.0001) (**Fig. 5C**). The failure to stimulate branching likely reflects that higher PGRN concentrations are required to induce sprouting of motor neurons as previously published^43^. In marked contrast to wt mice, *Sorcs2^-/-^* motor neurons were completely refractory to PGRN, as the total neurite length was identical to that of the unstimulated cultures (no stimulation: 156 μm, PGRN: 157 μm) (**Fig. 5E**). This was not accounted for by an intrinsic failure of the neurons to respond to trophic stimulation, since addition of BDNF increased both branching and neurite length in the *Sorcs2^-/-^* cultures (no stimulation: 2.14 branches, BDNF: 3.39 branches, p < 0.0001, and total length BDNF: 156 vs. 212 μm, p = 0.0001) (**Fig. 5D-E**). This was also seen for MNs from wt mice (2.16 branches vs. 3.46, p < 0.0001, and 156 μm vs. 246 μm, p < 0.0001) (**Fig. 5B-C**). Comparisons of the total branching and neurite length between genotypes showed that cultured motor neurons grew to a similar extent and responded equally well to BDNF treatment (branching: p = 0.997, length: p = 0.395), whereas the PGRN-induced increase in neurite length was significantly different between wt neurons compared to *Sorcs2^-/-^* (p = 0.0099) (**Fig. 5F-G**). We conclude that SorCS2 is required for PGRN-dependent motor neuron outgrowth *in vitro*.

**Figure 5:**
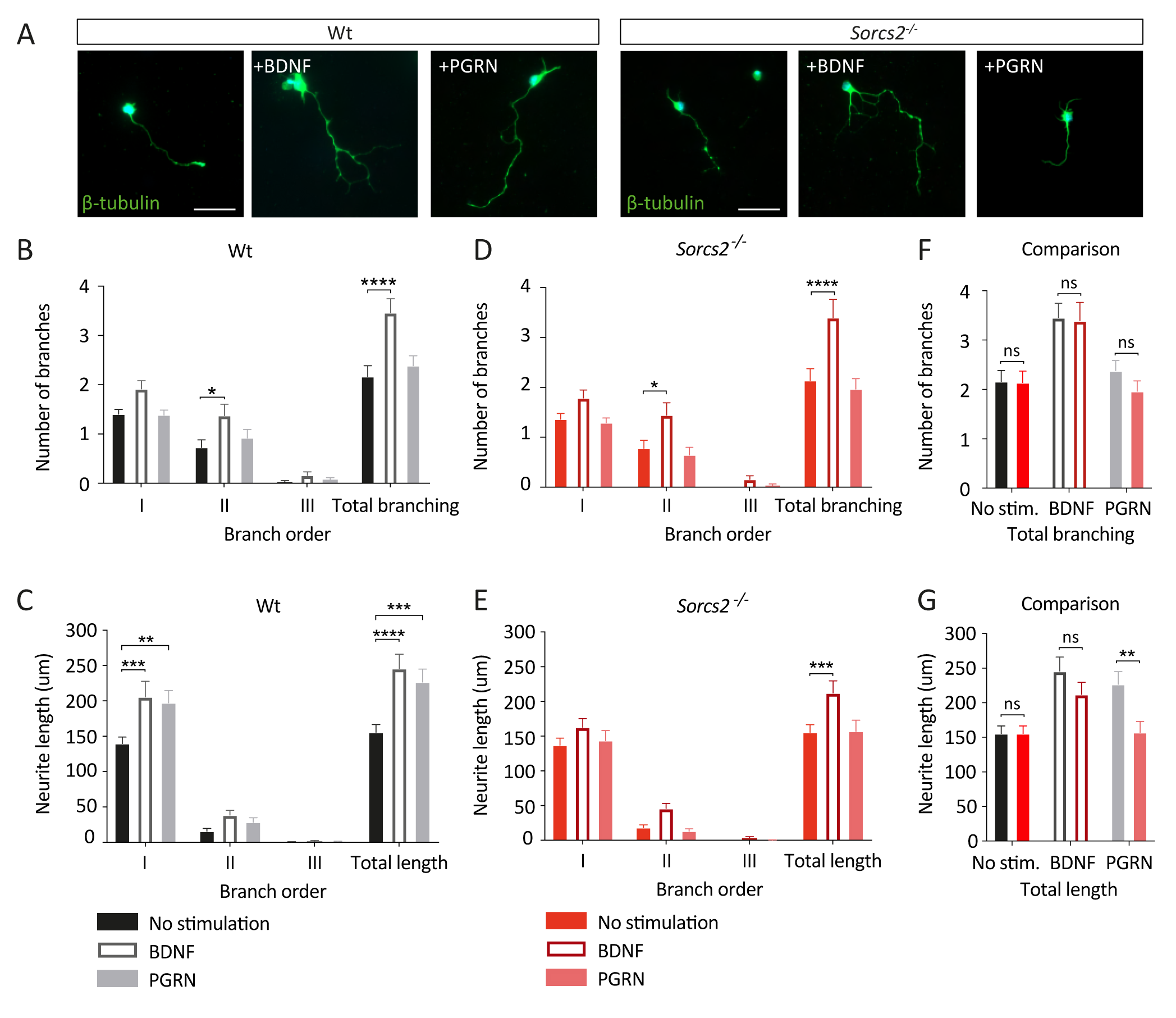
PGRN-induced motor neurite outgrowth requires SorCS2 expression. Primary motor neurons isolated from E13.5 murine spinal cord were cultured for 4 days. **(A)** Neurite outgrowth visualized by β-tubulin staining (green) in representative wt and *Sorcs2^-/-^* motor neuron cultures grown either without stimulation, with 5 ng/mL BDNF, or 100 ng/mL PGRN. Scalebar = 50 μm. Number of branches **(B+D)** and neurite length **(C+E)** of primary, secondary, tertiary neurites. The total neurite length or branching are summarized for wt (grey tones) and *Sorcs2^-/-^* (red tones) motor neurons (Wt; no stim. = 37 neurons, BDNF = 37, PGRN = 39, *sorcs2^-/-^*; no stim = 36, BDNF = 38, PGRN = 31). Comparisons of total branching **(F)** and total neurite length **(G)** between genotypes show that only PGRN stimulation of neurite outgrowth is significantly different between the two. Data shown as mean ± s.e.m. n = neurons. P-value determined with two-way ANOVA and Dunnet’s multiple comparisons test (C-E) or Šídák (F-G). * < 0.05, ** < 0.01, *** < 0.001, **** <0.0001.

### SorCS2 and PGRN are co-expressed in motor neurons postnatally

During postnatal development, the dendritic tree of the motor neurons grows to attain their maturity and to establish fully functioning synapses^44^. In the adult mouse PGRN supports neuronal function and is neuroprotective^33,45^. Hence, we explored SorCS2 and PGRN expression in motor neurons of the ventral horn in P3 (postnatal day 3) and adult mice 12 weeks of age. Double immunostaining for GFP and the motor neuron marker choline acetyltransferase (ChAT) in *Hb9::GFP* mice confirmed that Hb9 labels motor neurons also in the postnatal period (**Fig. 6A**). We found that SorCS2 and PGRN were abundant in *Hb9::GFP*-positive neurons, where they predominated in perinuclear vesicles in a pattern resembling that of the TGN (arrow heads, **Fig. 6B-C**). Furthermore, we extracted RNA sequencing data from laser-captured micro-dissected motor neurons^46^ to evaluate transcriptional profiles of *Sorcs2* and *Grn* in the spinal cord, the facial nucleus, and the cranial nuclei 3 and 4 in P2, P5, and P10 mice. The data showed that *Grn* and *Sorcs2* expression is not restricted to motor neurons originating from the spinal cord during the early postnatal period (**Fig. 6D**). In 12-weeks-old mice, RT-qPCR demonstrated that *Sorcs2* mRNA transcripts are prominent in the spinal cord as it is expressed to a level comparable to that in cortex (**Fig. 6E**). Protein expression was validated by immunoblotting (**Fig. 6F**), and IF microscopy of serial sections showed that in the spinal cord SorCS2 protein was restricted to ChAT+ motor neurons (**Fig. 6G-I**). Similarly, we detected PGRN in the adult spinal cord with expression being enriched in motor neurons (**Fig. S6A-C**); an observation that has also been reported by others^21^. Taken together, the data demonstrate that during early postnatal stages and in the adult mouse SorCS2 and PGRN are co-expressed in motor neurons suggesting a possible interaction inside the cell or at the cell surface following autocrine secretion.

**Figure 6:**
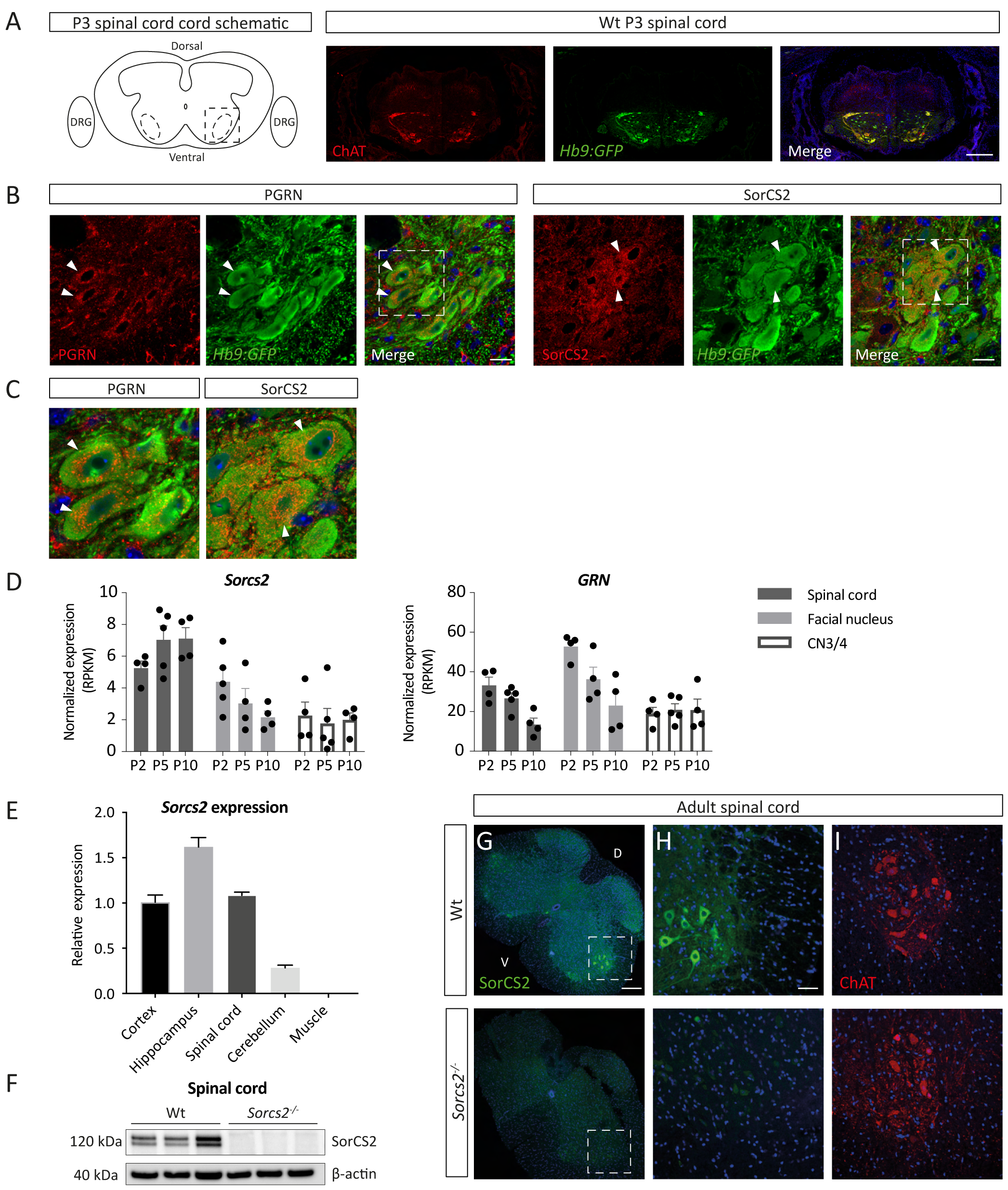
SorCS2 and PGRN are expressed in motor neurons postnatally. **(A-C)** Immunofluorescent staining of horizontal P3 spinal cord sections from *Hb9::GFP* mice. (A) Left, illustration of the section used. Right, ChAT (red) expression in GFP-labeled motor neurons confirming that Hb9 is also present in postmitotic motor neurons. Scalebar = 200 μm. **(B)** Expression of PGRN (left) and SorCS2 (right) in Hb9-positive motor neurons (arrow heads). Scalebar = 20 μm. Dashed squares are enlarged in **(C)**. Nuclei labeled with Hoechst in all sections. **(D)** Transcriptional profiling of laser-captured motor neurons from the spinal cord, the facial nucleus, and the cranial nuclei 3 and 4 from confirms *Grn* and *Sorcs2* expression in P2, P5, and P10 mice^46^. **(E)** RT-qPCR for *Sorcs2* expression in cortex, hippocampus, spinal cord, cerebellum and muscle in 12 weeks old wt mice (n = 3). Relative expression indicates fold-change normalized to cortical expression (2^-ΔΔCt^), shown as mean ± s.e.m. **(F)** Immunoblotting for SorCS2 in spinal cord demonstrates SorCS2 protein in Wt but not in *Sorcs2^-/-^* mice. β-actin serves as loading control. **(G)** Immunofluorescent staining of SorCS2 in adult murine wt and *Sorcs2^-/-^* spinal cord sections. Scalebar = 200 μm. **(H)** Enlargement of lateral ventral horn (white square). Motor neurons in wt but not *Sorcs2^-/-^* mice demonstrate SorCS2 expression (green). Scalebar = 50 μm. **(I)** Equivalent sections of ventral spinal cord showing ChAT expression (red) in motor neurons. Nuclei are stained using Hoechst.

### *Sorcs2^-/-^* mice show delayed functional recovery after facial nerve crush injury

PGRN was recently found to support axonal outgrowth in adult mice as *Grn^-/-^* mice showed delayed recovery after a facial nerve injury^27^. The facial nerve is made up entirely by motor axons originating from the facial motor nucleus in the brain stem that is required to move the whiskers (**Fig. 7A**). Double immunostaining with SMI-32 showed that SorCS2 was present in these motor neurons that previously have been reported also to express PGRN^27^ (**Fig. 7B**). We therefore investigated regeneration of the facial nerve in *Sorcs2^-/-^* mice following an injury by applying the paradigm used in the study on *Grn^-/-^* mice^27^. In this paradigm, the facial nerve is gently crushed distal to the retroauricular branch on one side, and the uninjured contralateral side is used as an internal control (**Fig. 7C**). After the crush, injured axons are lost and the whiskers become immobile. Regeneration is subsequently assessed by scoring provoked whisker movement twice a day. Whereas wt mice were fully recovered within 12 days, 14 days were required for the *Sorcs2^-/-^* mice (**Fig. 7D**). Quantified as the area under the curve of whisker movement, recovery was significantly delayed in the knockouts (p < 0.001) (**Fig. 7E**). Remarkably, the delayed recovery in the SorCS2 knockouts fully phenocopied that previously observed for *Grn^-/-^* mice^27^. Our data suggest that SorCS2 may control PGRN activity when co-expressed in the adult motor neurons during regeneration.

**Figure 7:**
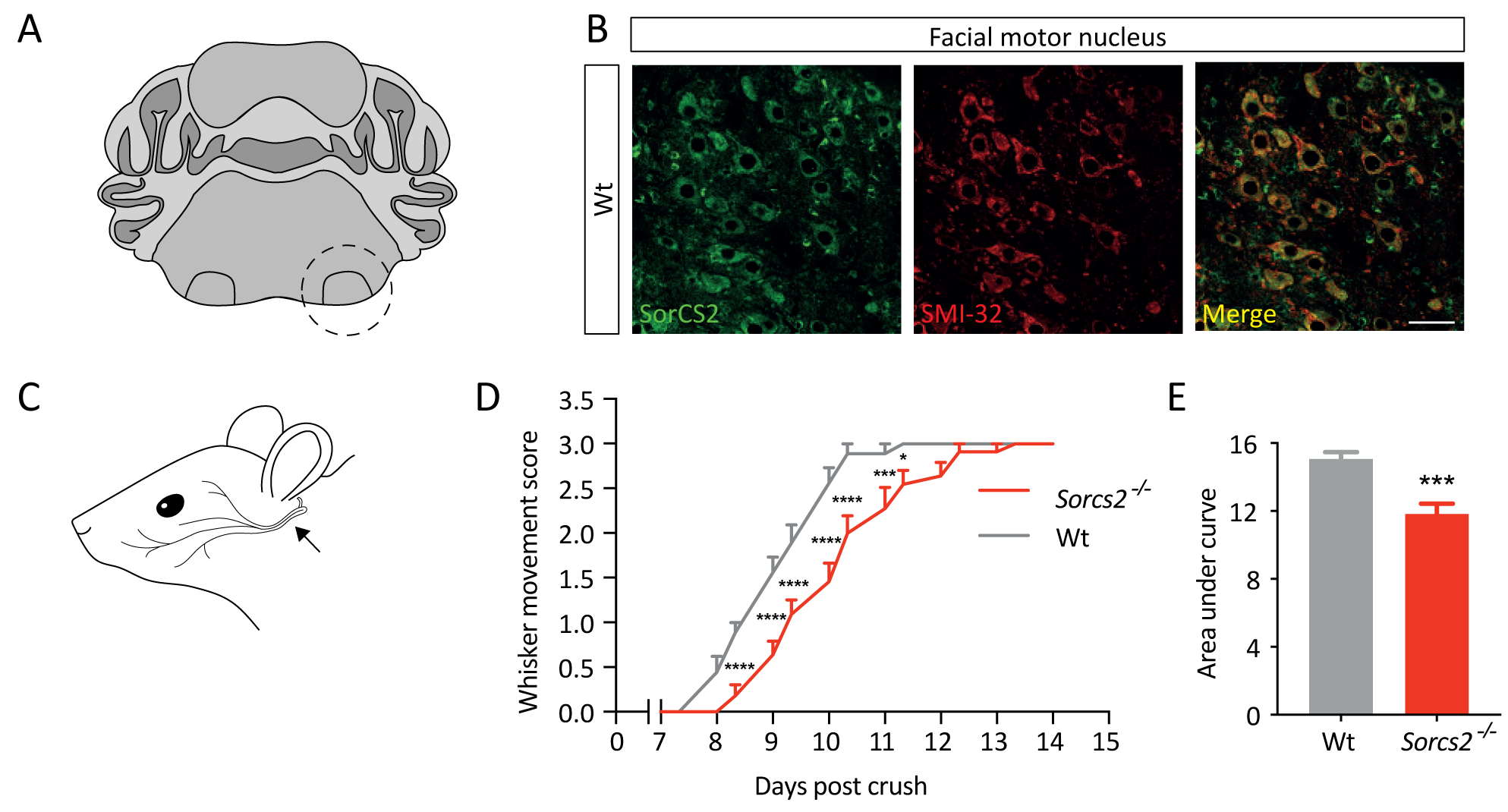
*Sorcs2^-/-^* mice show delayed functional recovery after facial nerve crush injury. **(A)** llustration of the brain stem with the facial motor nuclei outlined. **(B)** Immunofluorescent staining of SorCS2 (green) and the neuronal marker SMI-32 (red) in a section of the facial motor nucleus as outlined in panel (A) Scalebar = 50 μm. (**C**) Schematic representation of the facial nerve and the injury site. (**D**) Regeneration of the facial nerve scored based on whisker movement; 0: no movement, 1: detectable movement of individual whiskers, 2: significant but asymmetric motion, 3: symmetric voluntary motion. Lack of SorCS2 significantly delays functional recovery after injury (two-way ANOVA). (**E**) Area under the curve of whisker movement recovery scores is significantly reduced in *Sorcs2^-/-^* mice (student’s t-test) (n = 9,11). Data shown as mean ± s.e.m. **** < 0.0001, *** < 0.001, * < 0.05.

### SorCS2 binds PGRN to control its secretion

Given the direct binding of PRGN to SorCS2 and the requirement of SorCS2 for motor neurons to respond to exogenous PGRN by axon outgrowth (cf. **Figs. 4A and 5G**), we co-cultured HEK293 cells transfected with either PGRN or SorCS2, reasoning that the secreted PGRN would bind to neighbouring SorCS2-positive cells (*trans*-interaction). Surprisingly, we failed to detect any *trans*-interaction whether analysed by double IF microscopy (**Fig. 8A**) or by co-immunoprecipitation (co-IP) with anti-PGRN and probing the precipitate for SorCS2 (**Fig. 8C**). Similar results were obtained with N-terminally HA-tagged PGRN, which excludes the possibility that anti-PGRN antibody would occupy the binding site for SorCS2 (**Fig. S7A+C**). The result follows other studies demonstrating that at the plasma membrane, SorCS2 requires a co-receptor to efficiently bind its extracellular ligands^13^.

**Figure 8:**
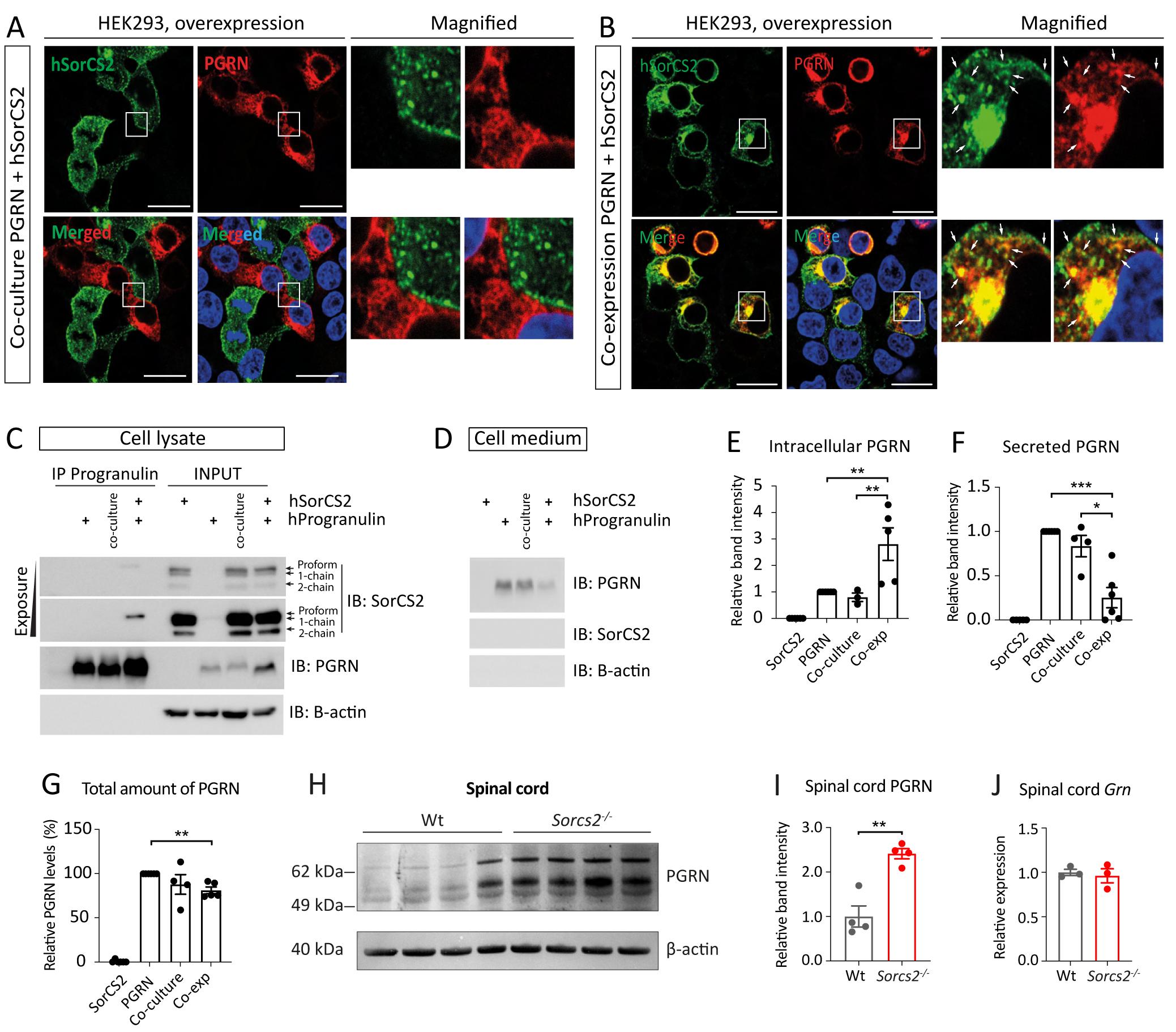
SorCS2 binds PGRN to control its secretion. **(A-B)** Immunofluorescence of HEK293 cells overexpressing human progranulin and SorCS2 separately and co-cultured **(A)** or co-expressing the receptors in the same cells (B). PGRN and SorCS2 do not colocalize on the plasma membrane when expressed separately (A), but they do so in Golgi-like compartments and vesicles when co-expressed in the same cells, highlighted by the arrows in (B). The nuclei are stained by Hoechst. Scalebar = 20 µm. **(C)** Co-immunoprecipitation of overexpressed SorCS2 and PGRN in HEK293 cells. SorCS2 physically binds PGRN intracellularly when co-expressed in the same cell (*cis*-configuration). The arrows point at three different isoforms of SorCS2 with the proform being accountable for the binding to PGRN. No co-IP is observed for the co-cultures **(D)** Representative immunoblots (IB) of the cell medium demonstrating secretion of PGRN. **(E-F)** In co-expressing cells, SorCS2 causes accumulation of PGRN intracellularly while reducing its secretion into the medium. The bars represent compiled quantification of the independent input samples blotted for PGRN (C+D) and HA in (S6C+D). The band intensity was measured as peak area (square pixel = 1) in ImageJ. **(G)** Total PGRN levels are decreased in cells co-expressing PGRN and SorCS2, quantified as combined relative band intensity from lysate and media (C+D). Immunoblots of β-actin serve as a loading control and negative control for the co-IP. P-value determined with Kruskal-Wallis test with uncorrected Dunn’s multiple comparison test **(E-G)**. **(H)** Immunoblotting for PGRN shows increased levels in spinal cord from P3 *Sorcs2^-/-^* mice. **(I)** Quantification of immunoblots using loading control for normalization. P-value < 0.01 (student’s t-test). **(J)** RT-qPCR of *Grn* expression in P3 spinal cord (n = 3). Relative expression indicates fold-change normalized to Wt (2^-ΔΔCt^). Data shown as mean ± s.e.m. *** < 0.001, ** < 0.01, * < 0.05.

Aside from signaling^13–15^, SorCS2 also mediates endocytosis and intracellular trafficking^16,18,47^. To assess whether SorCS2 and PGRN may bind in a *cis*-configuration, HEK293 cells were co-transfected to express both proteins simultaneously. We found that SorCS2 and PGRN strongly co-localized in a perinuclear compartment and to a lesser extent in vesicles dispersed throughout the cytoplasm, likely representing TGN, endosome and recycling vesicles (**Fig. 8B, S7B**). To demonstrate a physical interaction between the proteins in *cis*, PGRN was immunoprecipitated and the precipitate immunoblotted for SorCS2 (**Fig. 8C**). Remarkably, we found that PGRN bound SorCS2 and that this interaction was restricted to the proform of the receptor that is enriched in the TGN. A similar result was found when overexpressing HA-PGRN (**Fig. S6C**). Notably, overexpression of SorCS2 resulted in a *∼*300% increase in cell-associated PGRN (p = 0.0076) (**Fig. 8C+E, S7C**), that was paralleled by a corresponding decrease in PGRN secretion (p = 0.0010) (**Fig. 8D+F, S7D**). The total amount of PGRN produced by the cells, meaning the sum of PGRN in the medium and lysate, was decreased too, suggesting that SorCS2 may control PGRN secretion and potentially also its trafficking from the TGN to lysosomes for degradation or processing into granulins (**Fig. 8G**, p = 0.0068). To assess whether SorCS2 also alters PGRN levels when expressed at endogenous levels, we measured its expression in the spinal cord of *Sorcs2^-/-^* mice. In contrast to cells overexpressing SorCS2, PGRN in the spinal cord at P3 was increased by 240% in *Sorcs2^-/-^* mice (**Fig. 8H-I**, p = 0.0017) despite identical mRNA levels (**Fig. 8J**). An increase in PGRN was confirmed using another antibody and cohort of animals (49% increase, p = 0.015) (**Fig. S7E**). Overall, we conclude that SorCS2 can bind and retain PGRN in the biosynthetic pathway thereby regulating its secretion and extracellular concentration.

## Discussion

In this study, we identified SorCS2 as a novel regulator of motor axon outgrowth. Knockdown of *sorcs2* in zebrafish embryos resulted in delayed axon development and aberrant branching, defects in NMJ synapse formation, and ultimately in reduced motility. In *Sorcs2^-/-^* mice, innervation of the forelimbs and the upper trunk at E12.5 was substantially compromised as manifested by shorter axons and a reduced area covered by the terminal motor axon arbors. The functional recovery after a facial nerve injury was markedly slowed in the SorCS2-deficient mice too, which demonstrates a critical function of the receptor also in motor neuron regeneration.

We and others previously showed that SorCS2 regulates axonal innervation in the central nervous system, as binding of proBDNF or proNGF to SorCS2 in complex with p75^NTR^ causes growth cone collapse of CNS neurons^13–15^. As a consequence, *Sorcs2^-/-^* mice exhibit dopaminergic hyperinnervation of the prefrontal cortex^13^. Here we found that in motor neurons, SorCS2 enables axon outgrowth rather than its retraction, suggesting SorCS2 may induce opposing effects in distinct types of neurons. In support of neuron differentiated functions, primary motor neurons from the *Sorcs2^-/-^* mice responded to BDNF by axonal outgrowth and branching, whereas hippocampal neuron cultures have been shown not to do so^16^.

From a panel of proteins implicated in motor neuron development, we found robust binding of PGRN to SorCS2. The activities of PGRN are multiple and several receptors have been suggested. When bound to TNF receptor I/II, Toll-like receptor 9 and EphA2, PGRN may elicit anti-inflammatory effects and modulate innate immunity and angiogenesis, respectively^48–50^. The mechanisms underlying the trophic responses are less well understood. Notably however, it has been demonstrated that PGRN engagement of Notch receptors promote motor neuron survival and regeneration^51^. Sortilin, a paralogue of SorCS2 and the first PGRN receptor to be identified, efficiently binds and mediates endocytosis and lysosomal sorting of PGRN^31^ but the trophic effects of PGRN is independent of this receptor^43,52^. These observations are in marked contrast to those for SorCS2, as SorCS2 was required for PGRN to elicit its neurotrophic effects.

During development we found SorCS2 expression in motor neurons, whereas PGRN is abundant in cells in close proximity to the projecting axons. In E12.5 mice, the PGRN expressing cells were positive for Iba-1, corroborating previous reports that find PGRN is enriched in immune cells including macrophages and microglia of embryonic zebrafish and mice^53,54^. PGRN is also secreted by activated Schwann cells following sciatic nerve injury to promote regrowth of their neighboring neurons^55^. Interestingly, zebrafish Schwann cells are derived from neural crest cells that migrate along the motor axons^56^ correlating well with our observed *grna* expression pattern. Similar to microglia, neural crest cells can secrete factors to change their microenvironment^57^ and they may represent another source of PGRN. Taken together, the expression patterns suggest that PGRN released by Iba-1+ and Schwann cells, potentially also neural crest cells, may induce its neurotrophic effect on developing and regenerating motor neurons through binding to SorCS2.

Surprisingly, we failed to demonstrate *trans*-binding in transfected HEK293 cells, but various possibilities may explain this apparent paradox between lack of binding and biological activity. PGRN can be cleaved into individual granulins, GRN A-G, upon secretion^58^. The neurotrophic effect of PGRN is reliant on GRN E, as GRN E alone can induce survival and neuronal outgrowth, whereas PGRN without the GRN E domain lacks this capacity^20,52^. Previous studies demonstrated that GRN E is responsible for PGRN binding to sortilin suggesting the same may also apply for SorCS2^31^. We used antibodies directed against full-length PGRN and we cannot discount the possibility that GRN E is detected less efficiently, or that binding to SorCS2 sterically hinders subsequent antibody binding. This may also explain why we did not detect binding with the N-terminally HA-tagged PGRN when probed with the anti-HA antibody. A more intriguing explanation, however, is the requirement of a co-receptor for SorCS2. Conceptually, Vps10p-d receptors partner with co-receptors to form composite high affinity ligand binding sites^5,59^. For instance, the affinity of proNGF to sortilin is increased by 20-fold when in complex with p75^NTR 12^, and binding of proBDNF to SorCS2 or p75^NTR^ individually is barely detectable but when co-expressed surface labeling is greatly enhanced^13^. Likewise, it is possible that in addition to SorCS2, a co-receptor is required for PGRN to efficiently bind to the cell surface. This co-receptor may be absent from the SorCS2-transfected HEK293 cells, whereas in cultured motor neurons that respond to PGRN by neurite outgrowth, it is present. The identity of said co-receptor remains to be established but given PGRN can bind Notch receptors to stimulate nerve regeneration, it makes them good candidates^51^.

At E12.5 and in adult mice, we found that PGRN is also expressed by motor neurons. In these neurons and in transfected HEK293 cells that co-express SorCS2 and PGRN, the two proteins co-localized in structures compatible with TGN and sorting vesicles. Interestingly, by immunoprecipitation experiments we found that specifically the proform of SorCS2, which is enriched in the TGN, could bind PGRN in a *cis*-configuration and prevent its secretion. As total PGRN levels were decreased too when co-expressed in the HEK293 cells and PGRN was increased in spinal cords from *Sorcs2^-/-^* mice, it is possible that SorCS2 may also traffic PGRN from the TGN to lysosomes for degradation or processing into granulins. Notably, a similar complexity in function has also been reported for sortilin. At the plasma membrane sortilin enables proNGF/proBDNF signaling, whereas in the TGN it controls trafficking of BDNF to the regulated secretory pathway as well as to lysosomes for degradation^12,60–62^. Future studies should address the cell biological mechanisms by which SorCS2 controls PGRN sorting and signaling from the plasma membrane.

In conclusion we have identified SorCS2 as a novel PGRN receptor required for motor neuron outgrowth *in vivo* and *in vitro*. At the plasma membrane, SorCS2 enables PGRN signaling whereas inside the cell it controls PGRN secretion and thereby its extracellular bioavailability. SorCS2 is expressed in several neuronal subtypes and it is intriguing to speculate that the receptor is used more broadly to transduce trophic signals by PGRN and sustain neuronal integrity. Notably, SorCS2 is abundant in the cortical pyramidal neurons that degenerate in patients with frontotemporal dementia caused by *GRN* haploinsufficiency. We propose SorCS2 as a candidate target in development of disease-modifying treatments for motor neuron diseases and potentially for other neurodegenerative disorders.

## Materials & methods

### Zebrafish husbandry

Zebrafish were maintained according to institutional and national guidelines under breeding permit, 2017-15-0202-00098, from the Danish Animal Experiments Inspectorate under the Ministry of Justice. Zebrafish were kept at the facility at Molecular Biology and Genetics at Aarhus University under standard conditions with feeding four times a day on a 14 h light/10 h dark cycle at 28.5 °C. The AB strain obtained from the European Zebrafish resource Center was used to produce wildtype embryos for microinjections, RNA isolation, IHC and ISH. Zebrafish with the mutant allele *sorcs2*^sa11642^ with a nonsense mutation in *sorcs2* exon 10 was obtained from the Zebrafish Mutation Project^63^. Genotyping of mutant fish was performed by isolating genomic DNA from adult fish by swabbing^64^ and running PCR using KOD HotStart polymerase (Novagen) following standard protocol with 100 ng of DNA, 5% DMSO and primers 5’-TATGCACACCCTGATCTTATGG-3’ and 5’-CACACATACCTGGCTGTTGTTT-3’. Sequencing of PCR products at Eurofins Genomics was used to detect the specific point mutation in the mutant line. For live-imaging and *in situ* HCR the transgenic line *Tg(mnx1:GFP)^ml2^* from Zebrafish International Resource Center expressing GFP in the primary motor neurons was used. Embryos were collected after mating, staged by hpf^65^, and kept in E3 buffer (5 mM NaCl, 0.17 mM KCl, 0.33 mM CaCl_2_, 0.33 mM MgSO_4_, 0.001‰ methylene blue, 2 mM Hepes, pH 7.0) at 28°C.

### Zebrafish microinjections

A *sorcs2* targeted splice site-inhibiting MO (sMO) targeting the exon 2/intron 2 boundary (AAATCTTCTGTCACTTACCTCCACA) and standard control MO (cMO) were obtained from GeneTools, LLC. Given an injection volume of 5 nL, injection mixes were made with phenol red (0.2% in 1 mM HEPES, pH7.5), 5 ng of p53 MO to avoid unspecific apoptotic effects^66^, and either 5 ng of cMO or 3.7 ng of the sMO. For rescue experiments 500 pg/injection of human *SORCS2* mRNA was injected together with sMO. Injections were performed under a microscope using a pico-injector in embryos from the 1 to 4-cell stage. Human *SORCS2* mRNA was transcribed from a pcDNA3.1/zeo-SorCS2A-P716 plasmid (provided by Peder S. Madsen, Aarhus Univeristy) with mMessage mMachine T7 Transcription kit (Thermo Fisher) and purified with RNA MinElute kit (Qiagen).

### RT-PCR

RNA was purified from 30-50 embryos using RNeasy Mini Kit (Qiagen) and RT-PCR was performed to detect *sorcs2* mRNA in wt embryos using the Titanium One-Step RT-PCR kit (Clontech) with primers 5’-TTTTGCGCACCTGTACCCAGCTG-3’ and 5’-TAACGCGCTCCTGAAGCAGAGTC-3’ for detection of mRNA after injection of different amounts of sMO and 5’-TTTTGCGCACCTGTACCCAGCTGTGTT-3’ and 5’ TAACGCGCTCCTGAAGCAGAGTCCATT-3’ for the different developmental stages. As a reference, the *rpl13a* gene was used with primers 5’-TCTGGAGGACTGTAAGAGGTATGC-3’ and 5’-AGACGCACAATCTTGAGAGCAG-3’.

### Whole-mount zebrafish *in situ* hybridization

Whole-mount *in situ* hybridization was carried out as previously described in the 2010-updated version^67^ with addition of 5% Dextran Sulphate sodium salt (Sigma) to the hybridization mixture^68^. The *sorcs2* probe was transcribed using the DIG RNA Labelling Kit (Roche) with a T7 polymerase from a PCR fragment made with the following primers on embryonic cDNA (the T7 promotor sequence is shown in bold): 5’-GAAGACGCTCATCATGGCAG-3’ and 5’-**TAATACGACTCACTATAGGG**AATGAGGATGACGCTGCTGT-3’. Stained embryos were stored in 75% glycerol and images acquired using an Olympus IX71 microscope with an Olympus DP71 camera. Horizontal sections were obtained by molding stained embryos in Tissue-TEK O.C.T. (Sakura) on dry ice and sectioning on a Microm HM 560 cryostat.

### Whole-mount zebrafish immunohistochemistry

Whole-mount IHC was performed according to the protocol by GeneTex (www.genetex.com) with minor modifications. Zebrafish embryos of the specified stage were dechorionated using sharp forceps and fixated in 4% PFA ON at 4°C. Embryos were dehydrated and permeabilized according to protocol, but blocked with PBS + 0.1% Tween20 + 10% goat serum 1h at 4°C. They were incubated with znp-1 (1,25 μg/mL) in 0.1M potassium phosphate buffer (80% K_2_HPO_4_, 20% KH_2_PO_4_ + 0.8% Triton X100) at 4°C 16-72h. Embryos were incubated with goat-anti-mouse 488 (1:1000, Thermo Fisher) at RT 3h, immersed in 75% glycerol, and mounted laterally after removing the yolk sac^69^. They were imaged on a confocal laser-scanning microscope LSM-780 (Zeiss). For the motor axon analysis, the 20X objective was used to obtain z-stacks with step intervals of less than 2.3 μm of motor axons ranging from axon 5-16 counting rostrally. For analysis, each z-stack was split into the two hemi-segments of the axon rows and z-projected using ImageJ software^70^. The axons were scored using definitions described in ^71^ for length and ^72^ for branching.

For double staining of motor axons and AChRs, znp-1 and *α*-Bungarotoxin (BTX) were used with a modified protocol from Bio-protocol^73^. After ON fixation, embryos were washed with PBS and digested with 0.1% collagenase (1 mg/mL, Sigma) diluted in PBS with Ca^2+^ at RT 9 min for 27 hpf embryos and 45 min for 48 hpf. Embryos were incubated 30 min at RT with *α*-BTX-555 (10 μg/mL, Thermo Fisher) in PBT + 0.5% Triton X-100 + 10% NGS + 1% DMSO. They were then incubated with znp-1 antibody and secondary antibody as described.

### Whole-mount zebrafish third generation *in situ* hybridization chain reaction

*In situ* HCR v3.0^34^ was performed with probes and reagents provided by Molecular Instruments following their protocol. Zebrafish embryos were collected at 24, 27, and 48 hpf and fixated with 4% PFA ON at 4°C. 48 hpf embryos were furthermore bleached 1 h with 3% H_2_O_2_/0.5% KOH/PBS before dehydration to remove pigmentation. Following rehydration, the 48 hpf embryos were then treated with Proteinase K for 20 min at RT and postfixated 20 min RT. All embryos were incubated with 5 pmol/μl of probes ON at 37°C and hairpins as instructed. Two set of probes for *sorcs2* and *grna* were used to increase signal. After washing, embryos were mounted laterally with Prolong Glass Antifade Mountant (Thermo Fisher Scientific, P36984) after removing the yolk sac^69^. 10 μm cross sections were obtained of zebrafish trunk by molding stained embryos in O.C.T. on dry ice and sectioning on a Cryostar NX70. They were imaged on a confocal laser-scanning microscope LSM-780 (Zeiss). A 20X objective was used to obtain z-stacks with step intervals of 1.9 μm of trunk motor axons and cross sections. Z-stacks were z-projected for visualization using ImageJ software ensuring correct representation of co-localization with volume of projection listed in figure legends.

### Zebrafish live-imaging

cMO- and sMO-injected *Tg(mnx1:GFP)^ml2^* embryos were dechorionated and anesthetized with 0.05% tricaine in E3 buffer. They were laterally molded at 18 hpf into 0.5% low-melt agarose in an imaging dish with glass bottom and covered with E3 buffer containing tricaine to keep them sedated during imaging. Embryos were imaged from 19 hpf until 39 hpf with a Nikon T*i* Eclipse automated TIRF microscope equipped with a 10x differential interference contrast (DIC) air objective, a Perfect Focus 3 system and a Zyla sCMOS5.5 Megapixel camera (Andor) controlled by NIS Elements software from Nikon. The fluorescence illumination system was Cool LED-pE-300 white, and the fluorescence filter set used was standard GFP. Imaging, using widefield settings, was performed every 15 min for 20 h with 13 z-stacks separated with steps of 6 μm of both DIC light microscopy and fluorescence microscopy. Images and videos were analysed using NIS Elements and ImageJ software.

### Touch response assay for 48 hpf zebrafish embryos

48 hpf cMO and sMO embryos were dechorionated and returned to incubator for least an hour to avoid stress. In a petri dish with warmed E3, embryos were placed, one at a time, in the centre of concentric circles drawn with an increase in diameter of 0.5 cm and touched with an embryo poker^69^ at the back from behind. Using a Zeiss axio zoom V16 microscope with a Hamamatsu high speed camera, embryos were recorded for 10s collecting 100 images/s. Images were analysed using the Hokawo software (Hamamatsu) and time spent swimming 1 cm determined. Embryos that did not swim 1 cm or more were not included in analysis.

### List of primary antibodies

| Antibody | Source | IF | IF cells | Immunoblotting |
| --- | --- | --- | --- | --- |
| β-actin | A5441 Sigma, mouse | | | 1:5000 |
| ChAT | AB144P Merck Millipore, goat | 1:500 | 1:1000 |  |
| Foxp1 | Gift from J. Dasen | 1:1000 |  |  |
| GAPDH | Cell Signaling Technology |  |  | 1:3000 |
| GFP | Ab290 Abcam, rabbit | 1:1000 |  |  |
| HA | 18181 Abcam, mouse |  | 1:500 |  |
| HA | H6908 Sigma, rabbit |  |  | 1:500 |
| Hb9 | Gift from J. Dasen | 1:1600 |  |  |
| Iba-1 | #019-19741 Wako | 1:500 |  |  |
| Progranulin | AF2420 R&D, goat |  | 1:500 | 1:500 |
| Progranulin | MAB24201 R&D, mouse |  |  | 1:500 (Fig. 8H + S5) |
| Progranulin | AF2557 R&D, sheep | 1:250 |  | 1:1000 (Fig. S2E) |
| SM1-32<br>(neurofilament H) | Biolegend, mouse | 1:200 |  |  |
| SorCS2 | AF4237 R&D, sheep | 1:250<br>1:500 (E12.5) |  | 1:1000 |
| SorCS2 | F7100 Dako custom-made, rabbit |  | 1:2000 | 1:2000 |
| Tuj1 | MMS-435P Biosite, mouse |  | 1:1000 |  |

### Mouse background and breeding

Animal studies were conducted with C57BL/6j BomTac wt mice (Taconic), *Sorcs2^-/-^* mice^13^, and B6.Cg-Tg(Hlxb9-GFP)1Tmj/J (*Hb9::GFP)* mice (Jackson Laboratories). Animals were bred at the Animal Facility at Biomedicine Aarhus University, and experiments were approved by the Danish Animal Experiments Inspectorate under permit 2016-15-0202-00051 and carried out according to institutional and national guidelines. Animals were housed in the animal facility at Aarhus University with light-dark cycles of 12h with light from 6 a.m. to 6 p.m. They were fed standard chow and water *ad libitum* and their cages were cleaned every week and supplied with nesting, bedding material, and a tunnel.

### Section preparation and immunohistochemistry of mouse embryos

The protocol was kindly provided by Jeremy Dasen, NYU Langone, New York. E12.5 embryos were beheaded and eviscerated before being fixated in 4% PFA for 1h 30 min. Tissue was subsequently washed with PBS and kept in 30% sucrose ON at 4°C. Embryos were embedded in Tissue-TEK O.C.T. and 10 μm sections cut on a cryostat (Cryostar NX70). Please note that we used RNA scope sample preparation protocol prior to the IF of PGRN in E12.5 mouse embryos because it provided staining of higher quality in case of this specific antibody.

Slides were washed with PBS to remove O.C.T. They were blocked 20 min with PBS + 0.1% Triton X-100 + 1% BSA and incubated ON at 4°C with primary antibodies in blocking solution. Slides were washed with PBS + 0.1% Triton X-100 and incubated 60 min at RT with secondary antibodies (Jackson ImmunoResearch, Thermo Fisher) diluted 1:1000 in PBS + 0.1% Triton X-100. Mounted with Vectashield (Vector Lab) and imaged on a confocal laser-scanning microscope LSM 700 (Zeiss).

### Fluorescent RNA scope *in situ* hybridization

E12.5 mouse embryos were rinsed in PBS-pH7.4 (Invitrogen), and fixed in 4% PFA (Acros Organics) at 4°C for 24 hours. The tissue was gradually immersed in 10%, 20%, and 30% of sacharose/PBS-pH7.4 solution at 4°C until dehydrated. The embryos were washed, embedded in Tissue-TEK O.C.T. (Sakura), and frozen to −80°C. The tissue was cryo-sectioned (Cryostar NX70) as 14 µm slices that were air-dried in −20°C for 2 hours, and later stored at −80°C.

For the ISH, we implemented manufacturer instructions for fixed-frozen samples and RNAscope Multiplex Fluorescent Reagent Kit v2 kit (ACD Biotechne, #323100) using the following probes: Mm-Sorcs2-C1 (ACD Biotechne, #473411-C1) and Mn-Granulin-C3 (ACD Biotechne, #422861-C3). For the fluorescent labeling, we used Opal-690 reagent pack (Akoya Biosciences, FP1497001KT, 1:750) for the C1 probe, and Opal-570 reagent pack (Akoya Biosciences, #FP1488001KT, 1:2000) for the C3 probe. To secure good Hb9-GFP signal, we subsequently incubated the slides with rabbit anti-GFP antibody (Abcam, #ab290, 1:1000) in PBTA solution (0.3% Triton X-100, 0.1% BSA, 5% donkey serum) at 4°C ON. The next day, we used secondary antibody anti-rabbit Alexa-488 (Invitrogen, #A11008, 1:1000) together with Hoechst (Thermo Fisher Scientific, #33342, 1:2000) for 1h in RT. We mounted the slides with Prolong Gold Antifade Mountant (Thermo Fisher Scientific, *#*33342), and imaged them using confocal laser-scanning microscope Zeiss LSM 800.

### Section preparation and immunohistochemistry of postnatal day 3 and adult spinal cord

To preserve tissue morphology for IHC, transcardial perfusion fixation was performed on adult wt and *Sorcs2*^-/-^ mice, and spinal cords were dissected out and post-fixated in 4% PFA for 2-6 h. For P3, the spinal cords with surrounding vertebrates were dissected from *Hb9::GFP* mice and fixated in 4% PFA ON at 4°C. For both ages, the spinal cords were dehydrated in 30% sucrose ON at 4°C before freezing in Tissue-TEK O.C.T. on dry ice. 40 μm sections of adult tissue were cut on a Microm HM 560 cryostat and 10 μm sections of P3 on a Cryostar NX70, and they were stored at −80°C until further use.

For IHC staining slides were thawed and antigen-retrieval performed by incubating the sections with preheated Target Retrieval Solution (DAKO) 20 min at 70°C. Sections were incubated in Tris-buffered saline (TBS) + 5% donkey serum + 1% bovine serum albumin + 0.3% Triton X-100 for 30 min at RT for permeabilization of cells and blocking. Slides were incubated with primary antibodies ON at 4°C. The subsequent day, the sections were washed with TBS + 0.05% Triton X-100 and incubated with secondary antibodies (1:300, Thermo Fisher) 4h at RT. After incubation with Hoechst (0.2 μg/mL, Thermo), the slides were mounted using DAKO mounting medium. Images were acquired on a confocal laser-scanning microscope LSM-780 (Zeiss).

### Whole-mount immunostaining of mouse embryos

For whole-mount staining of embryonic motor nerves, heterozygous *Hb9::GFP* mice expressing GFP under the *Mnx* promotor were crossed with *Sorcs2^-/-^* or wt mice. The mice display GFP expression in dendrites, axons and soma of spinal motor neurons from E9.5 to P10. Embryos were collected at E12.5 and stained as demonstrated in ^74^. In brief, embryos were beheaded, eviscerated and pinned down on a silicone plate on ice with limbs slightly stretched out. Embryos were fixated ON at 4°C with 4% PFA. The embryos were then bleached 4h at 4°C with Dent’s bleach (1 part H_2_O_2_ : 2 parts Dent’s fix) followed by ON-fix with Dent’s fix (1 part DMSO : 4 parts MeOH). After washing with PBS, embryos were incubated in blocking solution consisting of PBS + 0.5% BSA + 20% DMSO 30 min at RT. Embryos were transferred to anti-GFP diluted in blocking solution 3 days ON gently rocking at RT. Embryos were then washed with PBS and incubated with donkey-anti-rabbit-488 (1:1000, Thermo Fisher) in blocking solution ON at RT. Following washes with PBS and MeOH, embryos were cleared with BABB (1 part benzyl alcohol : 2 parts benzyl benzoate). GFP-labelled motor axons were visualised in confocal z-stacks (around 400 μm thick) using a confocal laser-scanning microscope LSM-780 (Zeiss) and analysed using ImageJ software.

### Surface plasmon resonance analysis

Surface plasmon resonance was performed to assess protein-protein interaction and determine binding affinity on a Biacore 3000 instrument. Soluble SorCS2 was immobilized at a density of 0.0862 pmol/mm^2^ on a CM5 sensor chip. hPGRN was dialysed against 10 mM HEPES (pH 7.4), 150 mM NaCl, 1.5 mM CaCl_2_, 1 mM EGTA, and 0.005% surfactant P20 (running buffer), and 25–250 nM of the ligand was injected (5 ml/min) through the flow cell at 20°C. After 600 s, running buffer containing PGRN was substituted with buffer lacking the ligand, allowing for determination of both the association and dissociation constants. The binding response is expressed in relative response units, i.e., the difference in response between the immobilized protein flow cell and a corresponding control flow cell. Kinetic parameters were determined using the BIAevaluation 3.0 software.

### Primary motor neuron cultures

Primary motor neurons were prepared from E13.5 mice embryos as described in ^42^ with minor modifications. Panning plates were coated with monoclonal anti-murine p75^NTR^ antibody (1,46 μg/mL, Ab8877 Abcam) ON at 4°C diluted in sterile 10 mM Tris-HCl pH 9.5. Isolated motor neurons were counted with Neubauer counting chambers and seeded on Poly-DL-ornithine hydrobromide-(PORN, Sigma) and Laminin (Invitrogen)-coated coverslips in complete medium (Neurobasal medium supplemented with GlutaMax I, senicillin/streptomycin (1:100, Gibco), B27 (2%, Invitrogen), 2% horse serum). Cells were cultured at 5% CO2 and 37°C. On day 1 and 3, 80% of complete medium was changed to either new complete medium without trophic factors or complete medium containing 5 ng/mL BDNF (GF029 Merck Millipore) or 100 ng/mL human PGRN (AG-40A-0188Y Adipogen). On day 4, the cultures on coverslips were fixated with 4% PFA for 20 min, washed and stored in PBS at 4°C until immunostaining.

### Immunostaining of primary motor neurons and analysis

For immunostaining of the primary motor neurons, coverslips with neurons were permeabilized in PBS + 0.1% Triton X-100, blocked with PBS + 10% fetal bovine serum (FBS) 30 min at RT, and incubated with primary antibodies ChAT or Tuj1 (neuronal class III β-tubulin) diluted in blocking solution ON at 4°C. After washing, coverslips were incubated with secondary antibodies 2h at RT and nuclei stained with Hoechst (0.2 μg/mL). Coverslips were mounted with DAKO mounting medium and imaged using an Apotome 2.0 microscope (Zeiss). Neurons were measured using ImageJ, and neurites longer than twice the length of the soma and projecting directly from the cell body were considered primary. Secondary neurites were defined as neurites protruding from the primary, and tertiary as neurites protruding from the secondary. 31-39 neurons of each genotype and stimulation were analyzed from two independent experiments.

### Mouse RT-qPCR

Cortex, hippocampus, cerebellum, spinal cord and gastrocnemius muscle from wt and *Sorcs2^-/-^* mice were collected in RNAlater (Invitrogen) and left ON at 4°C. The following day the tissue was lysed in RLT buffer and RNA purified using RNeasy (Qiagen). cDNA was synthesized using the High Capacity RNA-to-cDNA kit (Thermo Fisher). RT-qPCR was performed using TaqMan Fast Advanced Master Mix (Applied Biosystems), TaqMan expression probes, and a 7500 Fast Real-Time PCR system (Thermo Fisher). *Hprt* and *Gapdh*, were used for normalization measuring fold change by 2^-ΔΔCt^, and three technical and three biological replicates were analyzed. All TaqMan probes were determined to be within 90-110% efficiency. TaqMan probes used (Thermo Fisher): *Sorcs2* (Mm00473050_m1), *Grn* (Mm00433848_m1), *Gapdh* (Mm99999915_g1), *Hprt* (Mm00446968_m1).

### Immunoblotting

Spinal cords from wt and *Sorcs2^-/-^* mice were dissected and lysed on ice with TNE lysis buffer (Tris-base 10 mM, NaCl 150 mM, EDTA 1 mM, NP40 1%, cOmplete 1% (Roche), PhosSTOP 1% (Roche)) and mechanically with a pellet pestle. Lysed tissue was centrifuged at 13,000 rpm and protein concentration of the supernatant measured using a bicinchoninic acid assay. Protein samples were denatured by boiling with SDS sample buffer and 20 mM DTT. Proteins were then separated by SDS-PAGE using a precast NuPAGE 4-12% Bis-Tris Gel (Invitrogen) and transferred to an PVDF membrane by immunoblotting in blotting buffer (192 mM glycine, 25 mM Tris-Base, pH 8.0). The membrane was subsequently blocked in 0.05 M Tris-base, 0.5 M NaCl, 0.1% Tween-20 (TST buffer) + 2% skimmed milk powder and 0.5% Tween-20 and washed in wash buffer (10 mM HEPES pH 7.8, 140 mM NaCl, 2 mM CaCl_2_, 1 mM mgCl_2_, 0.2% skimmed milk powder, 0.05% Tween-20). It was incubated with primary antibodies diluted in wash buffer for 2h at RT, then HRP-conjugated secondary antibodies (DAKO) and visualized with chemiluminescence using ECL Western Blotting Detection Reagents (GE Healthcare) and the imaging system FUJI Film LAS 4000 (GE Healthcare).

### Facial nerve crush injury

The facial nerve crush surgery was performed at VIB-KU Leuven and were approved by Institutional Animal Care and Ethical Research Advisory Committee of the University of Leuven, Belgium (P132/2012 and P015/2015). The surgeries were performed as described previously^27^ on 10 weeks old wt and *Sorcs2^-/-^* mice of both sexes that were shipped to Belgium from Denmark. To evaluate regeneration of facial nerve axons, all mice were examined twice daily (9-10 am and 6-7 pm) for whisker movement. Whisker movement scores were given according to the following scale: 0 for no detectable movement, 1 for detectable motion of individual whiskers, 2 for significant (but asymmetric) voluntary motion, and 3 for symmetric voluntary motion. The contralateral whiskers were used as a reference for complete functional recovery.

Immunostaining of the facial motor nucleus was performed as described previously^27^. Images were acquired using a Leica SP8x confocal microscope.

### Cell cultures, transfection and co-cultures

Human embryonic kidney 293 cells (HEK293, ATCC) were grown at 37°C with constant 95% of H_2_O and 5% of O_2_. Cells were cultivated in Dulbecco’s Modified Eagle’s medium (DMEM, Sigma) supplemented with 10% FBS and 50 U/ml penicillin/streptomycin (all Thermo). Cells were trypsinized in 1 ml of Trypsin_0.05%-EDTA (Thermo) and seeded into 10 cm dishes in 40% confluence 24 hours prior the experiment. The next day, the cells were transfected using FuGENE 6 transfection reagent (Promega). Transfection mixture was prepared in serum-reduced OptiMEM medium (Thermo) using 6 µg of plasmid DNA per dish (3 µg DNA1 : 3 µg DNA2). The following plasmid DNAs were used in this study: pcDNA3.1, pcDNA3.1-human_SorCS2_Full length, pCMV3-human_Progranulin (Sino Biological, #HG10826-UT) and pCMV3-HA-human_Progranulin (Sino Biological, # HG10826-NY). Transfection mixture was added to 7 ml of complete cell media and incubated for 24 hours. 24 hours post transfection, cells were trypsinized in 1 ml of trypsin and seeded into either 10 cm dishes for co-IP experiments (8 ml of fresh media plus 400 µl of cell suspension, or 400 µl + 400 µl of cell suspension for the co-cultures), or into 24-well plates that contained glass cover-slips coated with Poly-L-Lysine. For the coverslips, 3.5 ml of fresh media with 100 µl cell suspension was added for single-transfections, or 50 µl + 50 µl for the co cultures. 500 µl of this mixture was seeded per well. The cells were subsequently grown 24h and harvested 48h post transfection.

### Co-immunoprecipitation and immunoblotting

For the co-IP experiments, HEK293 cells were harvested either 24 or 48h post transfection. First, 2 ml of cell media was spun down at 200g for 5 min at RT to remove the cell debris. The samples were placed on ice and cells were washed with PBS, before adding 1 ml of TNE lysis buffer while shaking at 4°C for 20 min. Cell lysates and cell medium were spun down at 20,000g at 4°C for 15 min, and supernatant was kept for further processing. 60 µl from each sample was used as an INPUT reference sample and was mixed with 20 mM DTT and NuPAGE LDS sample buffer. 400-600 µl of the cell lysate per condition were used for the pulldown. The following antibodies were used: 1µg of progranulin (AF2420 R&D), 1.5 µg of rabbit anti-HA (H6908 Sigma), and 1 µg of mouse-anti-HA (26183 Sigma). Cell lysates were incubated with antibodies for 14-16h in 4°C. Meanwhile, 30 μl GammaBind G Sepharose beads (Sigma) per sample were washed with PBS, centrifuged at 300g for 5 min at RT, and washed in TNE lysis buffer before another centrifugation step at 300g for 5 min at RT. Washed beads were incubated with the antibody-lysate for 4h at 4°C with slow rotation. Samples were subsequently washed on ice with PBS + 0.1% Tween20 and spun down at 200g for 1 min at 4°C. Proteins were eluted by heat in elution buffer (TNE buffer, 20 mM DTT, NuPAGE LDS sample buffer). All the samples (including INPUT) were immediately boiled at 95°C for 5 minutes and subjected to immunoblotting using home-made 8% Tris-Glycin polyacrylamide SDS-Page gels.

Immunoblotting was performed as described in previous section. Secondary antibodies fused with HRP (DAKO) were used in dilution 1:2000. The immunoblots were developed dependent on the strength of the signal either with ECL Amersham RPN2106 (Sigma) or Immobilon Forte Western HRP substrate (Sigma). Images were processed using Adobe Photoshop CC. Band intensity was measured in ImageJ and analysed in Microsoft Excel. The band intensities of IB: PGRN and IB: HA were further normalized to the band intensity of *β*-actin (loading control). To describe relative band intensity, the values were further normalized to IB of Progranulin single transfection, which was set as 1.

### Immunostaining of HEK293 cells

Transfected HEK293 cells were harvested for immunofluorescence 48h post transfection. They were fixated with 4% PFA at RT for 15 min. Immunofluorescence was performed as described previously^75^. Secondary antibodies (Thermo) were used in dilution 1:800 for 1h at RT. The samples were imaged using scanning confocal microscopy system Zeiss LSM-780. The images were processed in Adobe Photoshop CC.

### Statistical analysis

Student’s T-test was used to compare differences in mean between two groups. Mann-Whitney test was used to compare ranks between non-normal distributed data. For multiple comparisons within the same experiment the Holm-Šídák method was used to correct for multiple comparisons with *α* =• 0.5 unless otherwise stated. For analysis of more than two groups, analysis of variance (ANOVA) was performed followed by Tukey’s multiple comparison test for comparing every mean with every other mean or Dunnett’s for comparing every mean with a control mean. Repeated measures two-way ANOVA with Šídák correction was used for the analysis of facial nerve recovery since values were matched according to time. All statistics was performed using the GraphPad Prism software. A p-value of < 0.05 was considered statistically significant. Data are presented as mean (standard error of mean, s.e.m.) unless otherwise stated.

### Analysis of deposited RNA sequencing data

The single-cell transcriptome atlas of zebrafish embryos^35^ on http://zebrafish-dev.cells.ucsc.edu was used to assess expression of *sorcs2* and *grna,*.

Transcriptional data from laser-captured motor neurons from the spinal cord, the facial nucleus, and the cranial nuclei 3 and 4^46^ was downloaded from NCBI Gene Expression Omnibus (GEO) (GSE115706), and normalized expression values (RPKM) extracted and plotted for *Sorcs2* and *Grn*.

## Acknowledgements

We wish to thank the entire Nykjaer laboratory for helpful discussions. Benedicte Vestergaard, Andreea-Cornelia Udrea, Anja Aagaard, and Anne Kerstine Thomassen are thanked for excellent technical assistance, and Peder S. Madsen for the SorCS2 plasmid. Thanks to Petra Kompanikova who helped establish the *in situ* HCR v3.0. PBT sends her gratitude to Moses Chao and his group for hosting her at NYU, and to Jeremy Dasen and his group members for all their help. The study was funded by the Lundbeck Foundation (Grant Nos. R90-2011-7723, R248-2017-431, and R315-2018-3066), The Danish National Research Foundation (Grant No. DNRF133), The Danish Council for Independent Research (Grant No. 7016-00261), The Bluefield Project (Grant No. BFP-AN-2022), EMBO (Grant No. 7375), and The Danish Society for Biochemistry and Molecular Biology. PVD holds a senior clinical investigatorship of FWO-Vlaanderen and is supported by the E. von Behring Chair for Neuromuscular and Neurodegenerative Disorders, the ALS Liga België and the KU Leuven funds “Een Hart voor ALS”, “Laeversfonds voor ALS Onderzoek” and the “Valéry Perrier Race against ALS Fund”.

## Author contributions

PBT performed the majority of experiments and data analysis and prepared the figures under supervision by AN. AS performed interaction studies (co-expression in HEK293 cells and IP) with help from PLO, analysis of murine RNA seq. data together with PQ, and IF of PGRN and RNA scope of *Grn* in E12.5 mice together with SN. AS helped prepare figures for the mentioned experiments. HL contributed to RT-PCR and prepared h*SORCS2* mRNA for rescue experiments. JTJ assisted with immunoblotting, in tissue collection, and in isolating, growing and staining primary motor neurons. SB performed the facial nerve crush and scored recovery together with PBT under supervision of PVD. JD provided mouse embryo sections, antibodies, and guidance for IF. Zebrafish studies were performed under supervision from KKS as well as HL, LNN, and CO. MVC contributed with scientific discussions and critical review of the manuscript. PBT, AS, and AN wrote the manuscript with input from all authors.

## Competing interests

The authors declare no competing interests.

---

**Figure S1:**
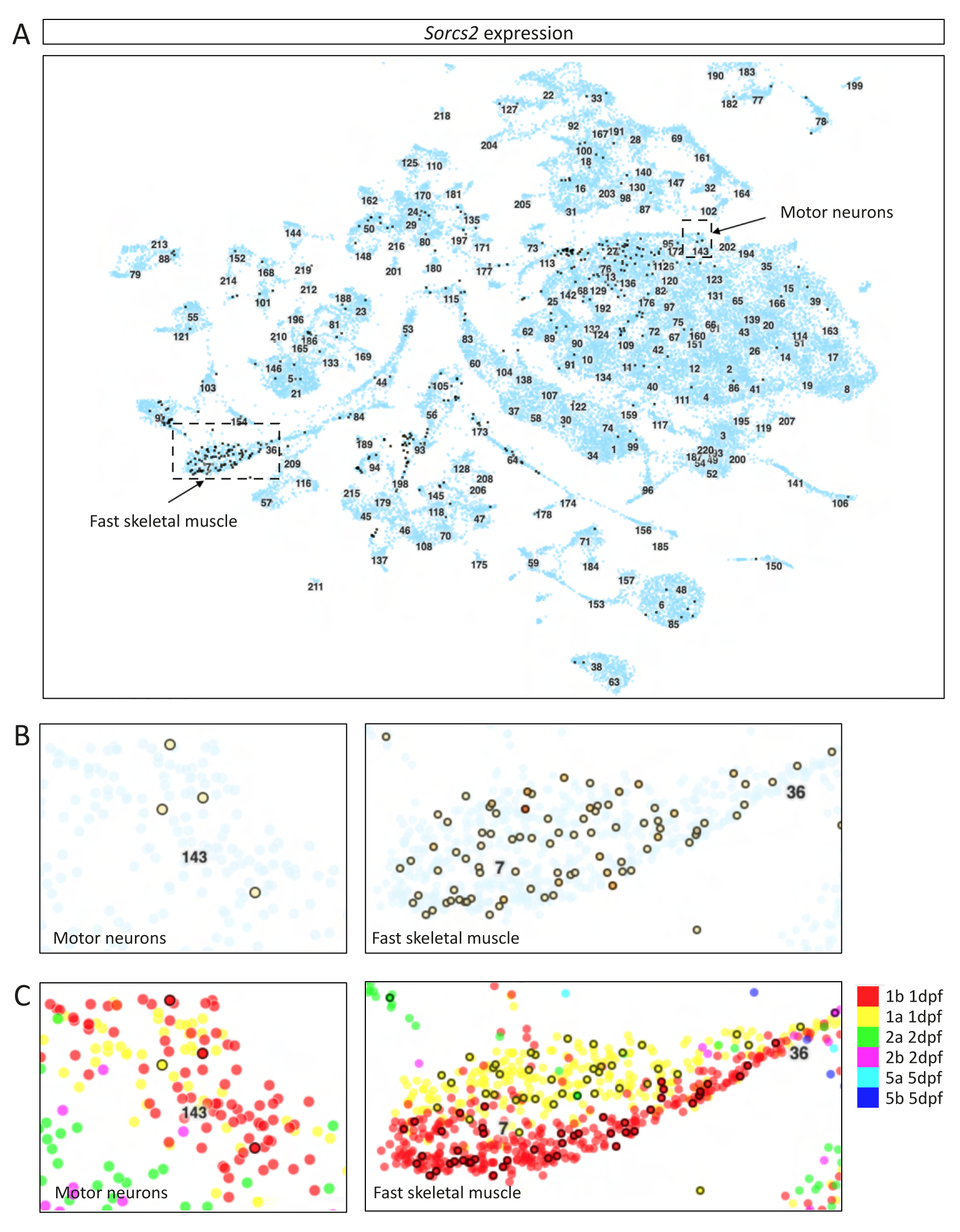
Single-cell transcriptome data of zebrafish *sorcs2*. Data from the single-cell transcriptome atlas of zebrafish embryos encompassing transcriptional profiles of 44,102 cells clustered according to cell type across four days of development^35^. **(A-C)** *Sorcs2* is expressed in different cell types including motor neurons and fast skeletal muscle in clusters 143, 7, and 36 (arrows). **(B)** Enlargement of dashed squares and with the age profile shown in **(C)**. dpf = days post fertilization.

**Figure S2:**
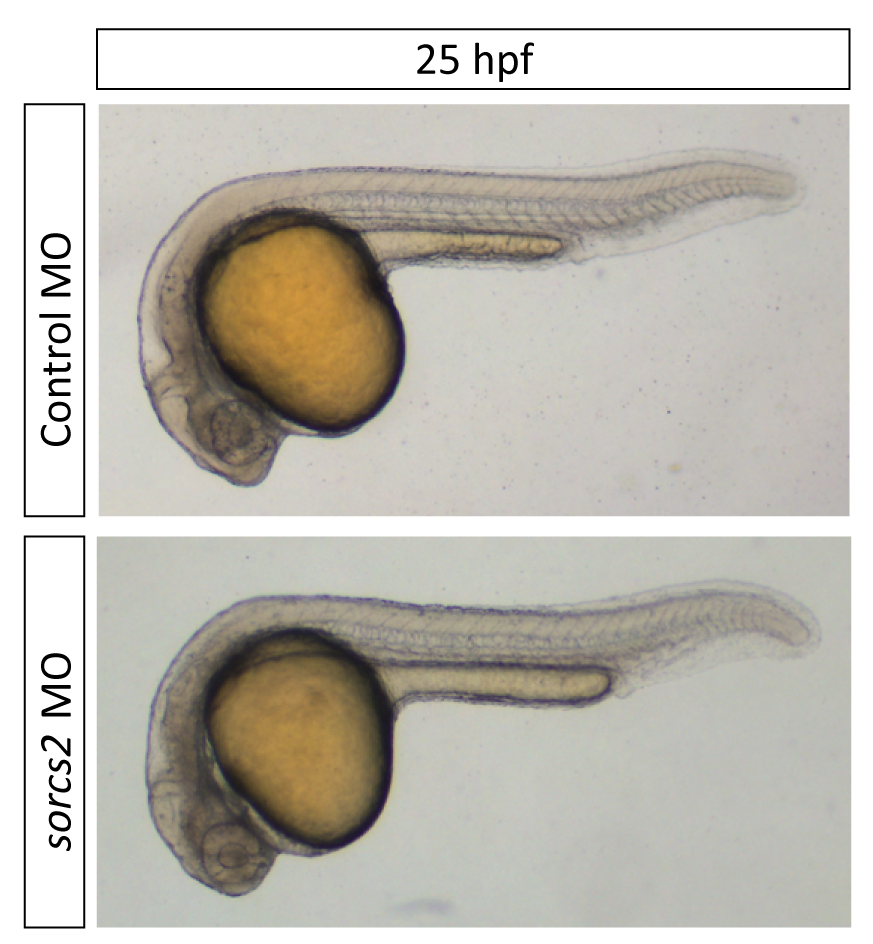
Bright field images of a control MO and a *sorcs2* MO embryo at 25 hpf.

**Figure S3:**
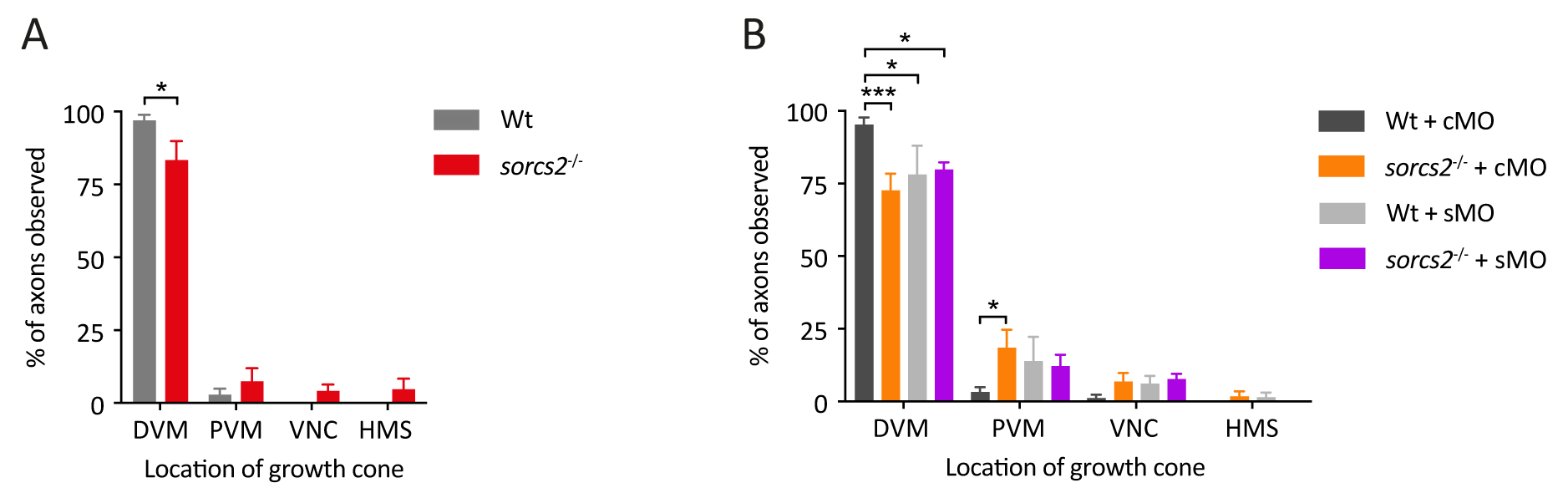
CaP axon outgrowth in *sorcs2^-/-^* zebrafish embryos. **(A-B)** Axon outgrowth in CaP axons from wt and *sorcs2^-/-^* embryos from heterozygous cross of *sorcs2*^sa11642^ zebrafish are shown in percentage. **(A)** *sorcs2^-/-^* axons (n = 6) are stunted compared to wt (n = 6) (p = 0.022). **(B)** Compared to wt + cMO (n = 6), axons from wt + sMO (n = 4, p = 0.011), *sorcs2^-/-^* + cMO (n = 4, p = 0.0005), and *sorcs2^-/-^* + sMO (n = 3, p = 0.047) are stunted. Injection of sMO into *sorcs2^-/-^* embryos does not aggravate the phenotype (73% vs. 80%, p = 0.670). Error bars: s.e.m., n = number of embryos. P-value determined with two-way ANOVA and Tukey’s multiple comparisons. *** < 0.001, * < 0.05. HMS, horizontal myoseptum; VNC, ventral edge of notochord; PVM and DVM, distal and proximal ventral myotome.

**Figure S4:**
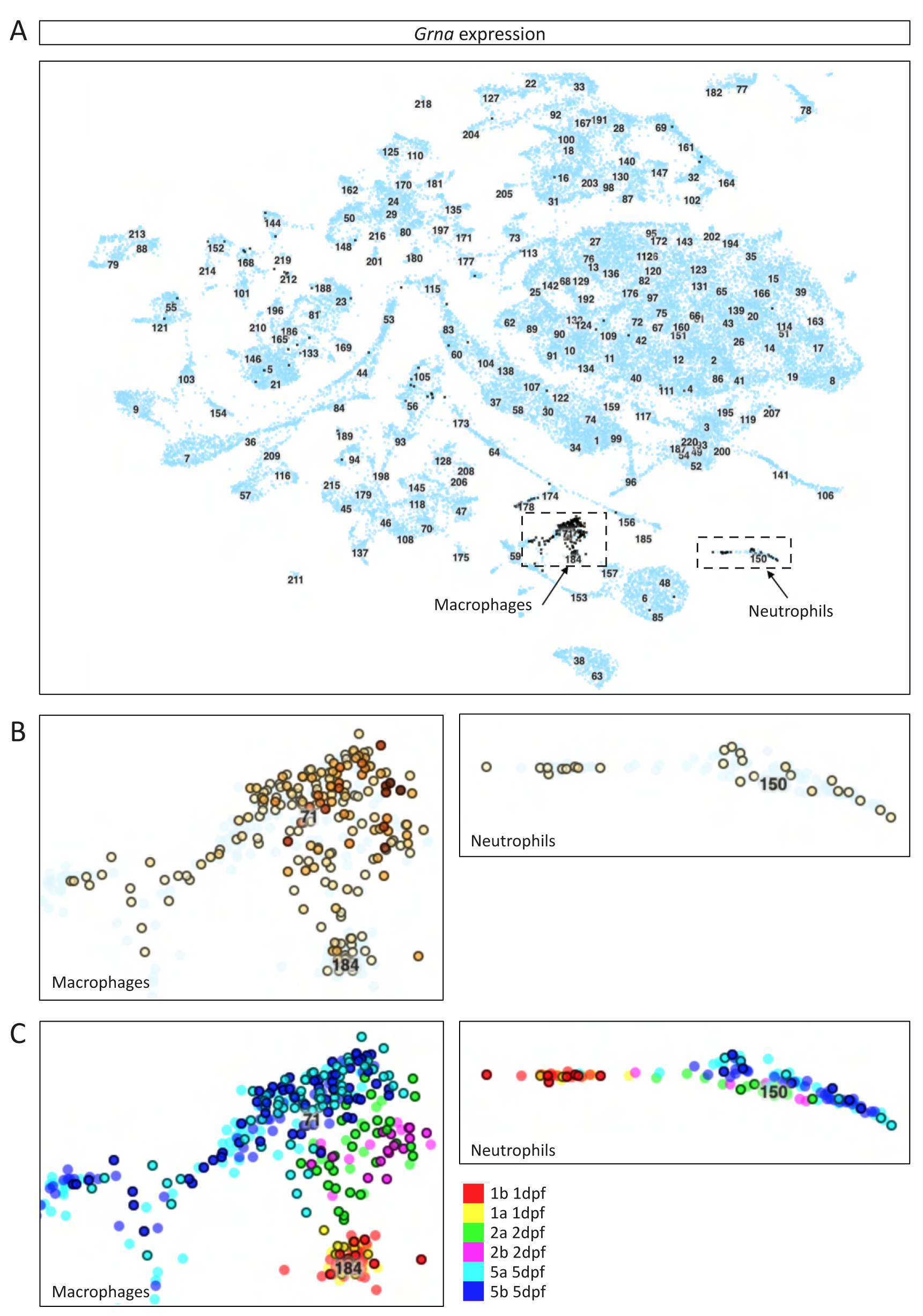
Single-cell transcriptome data of zebrafish *grna*. Data from the single-cell transcriptome atlas of zebrafish embryos encompassing transcriptional profiles of 44,102 cells clustered according to cell type across four days of development^35^. **(A-C)** *Grna* is mainly expressed in macrophages and neutrophils in clusters 71, 184, and 150 (arrows). **(B)** Enlargement of dashed squares and with the age profile shown in **(C)**. dpf = days post fertilization.

**Figure S5:**
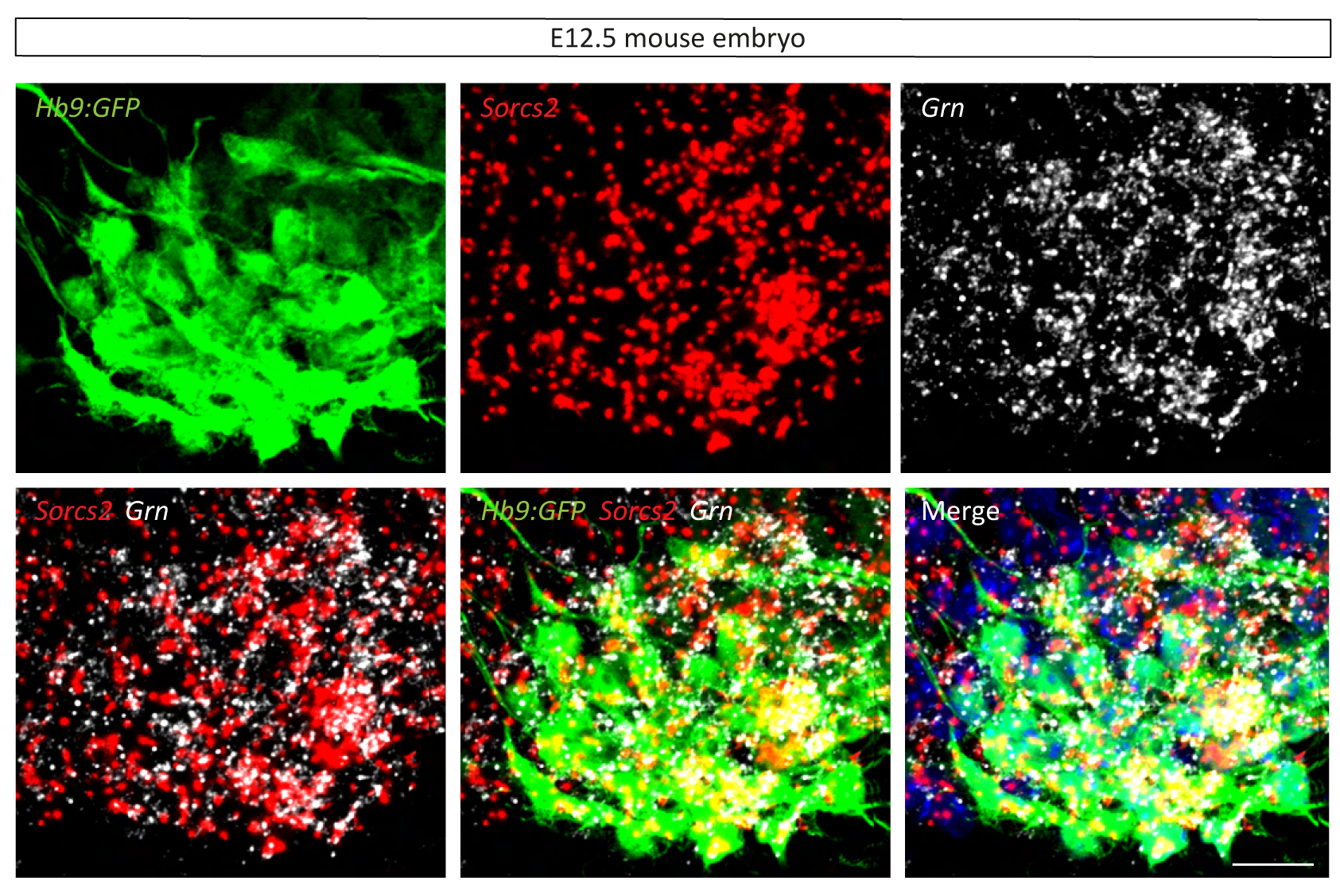
RNA scope *in situ* hybridization of *Sorcs2* and *Grn*. RNA scope *in situ* hybridization of *Sorcs2* (red) and *Grn* (white) in horizontal sections of E12.5 *Hb9::GFP* mice. Scalebar = 10 μm.

**Figure S6:**
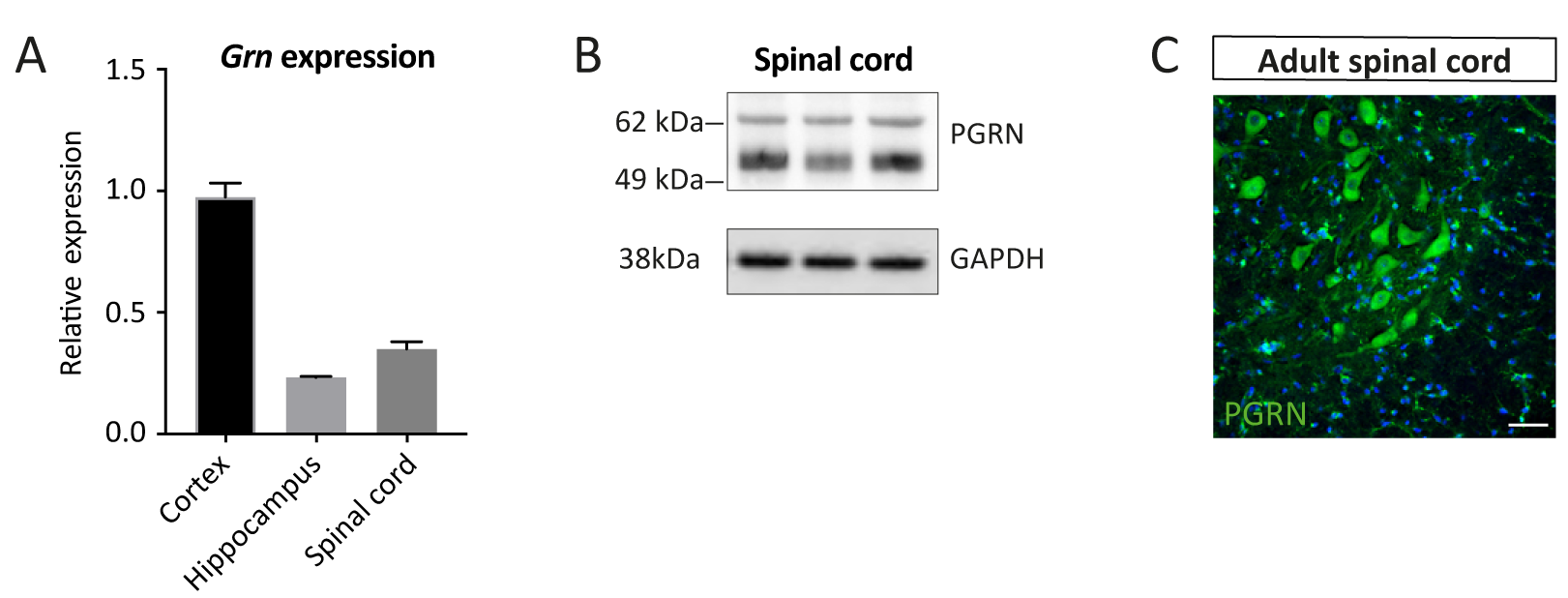
PGRN expression in adult murine spinal cord. **(A)** RT-qPCR for *Grn* expression in cortex, hippocampus, and spinal cord in 12 weeks old wt mice (n = 3). Relative expression indicates fold-change normalized to cortical expression (2^-ΔΔCt^), shown as mean ± s.e.m. **(F)** Immunoblotting for PGRN in spinal cord demonstrates PGRN protein in Wt mice. GAPDH serves as loading control. **(G)** Immunofluorescent staining of PGRN in the lateral ventral spinal cord from an adult mouse showing PGRN (green) in motor neurons. Scalebar = 50 μm. Nuclei are stained using Hoechst.

**Figure S7:**
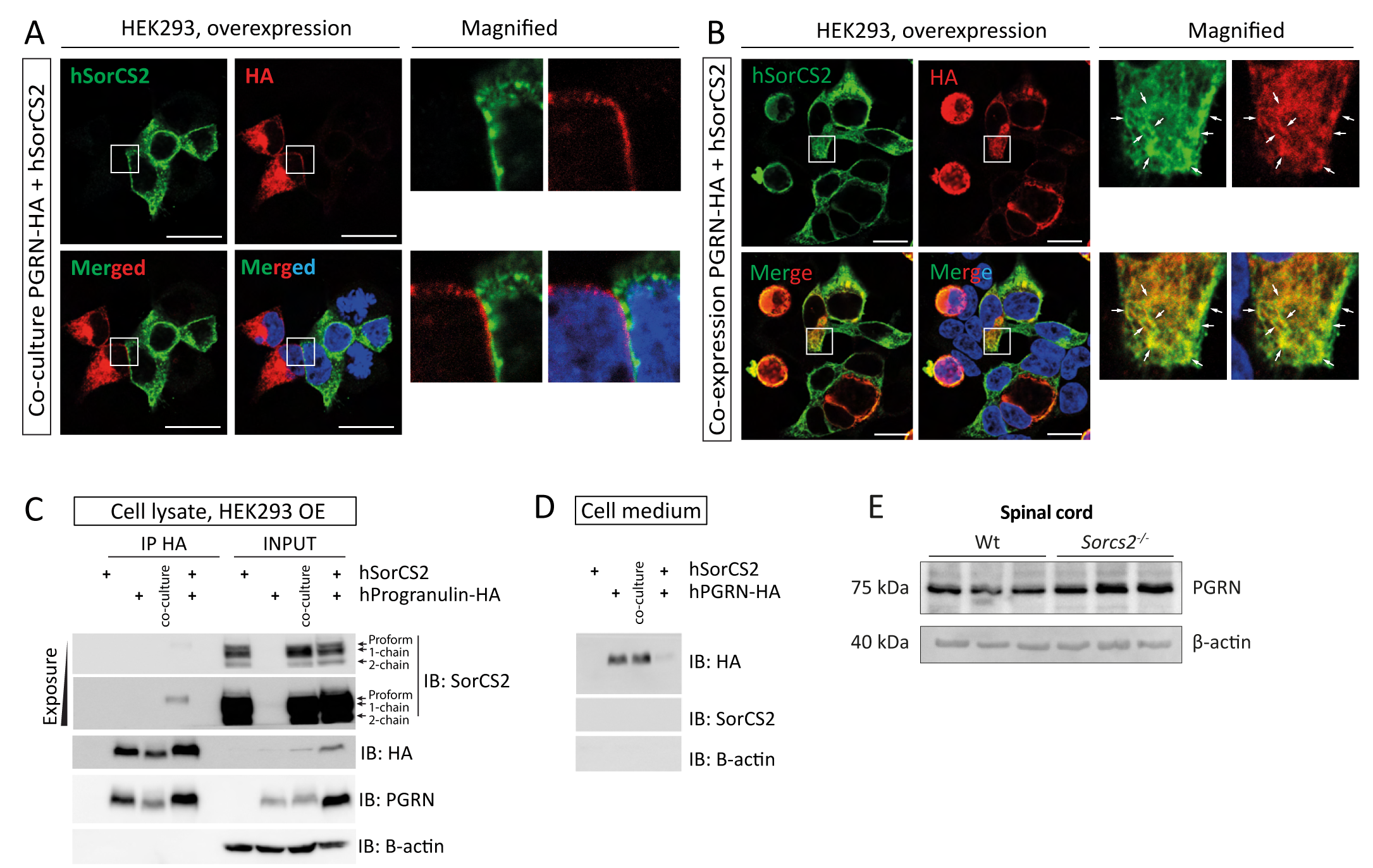
SorC2 and HA-PGRN binding and secretion. **(A-B)** Immunofluorescence of HEK293 cells overexpressing human HA-tagged PGRN (N-terminal tag) and SorCS2 separately and co-cultured (A) or co-expressing the receptors in the same cells (B). When expressed in *trans* (separately) there is no staining for PGRN at the plasma membrane in the SorCS2 expressing cells. (B) In *cis* (co-expression), SorCS2 co-localizes with HA-PGRN in Golgi-like compartments and vesicles as highlighted by the arrows in (B). The nuclei are stained using Hoechst. Scalebar = 20 µm. **(C)** Co-immunoprecipitation of SorCS2 with human HA-PGRN when co-expressed but not when expressed separately. The arrows point at three different isoforms of SorCS2 showing that the proform accounts for the binding to PGRN. **(D)** Immunoblots (IB) of the cell medium demonstrates secretion of PGRN. **(E)** Immunoblotting for PGRN shows that the levels in spinal cord from P3 *Sorcs2^-/-^* mice are increased. β-actin is used as loading control.

## Supplementary movies

**Supplementary movie 1:** Live-imaging of a *sorcs2* KD *Tg(mnx1:GFP)^ml2^* embryo from 18 to 38 hpf showing premature branching and truncation defects. Embryo oriented anterior to the right, dorsal to the top.

**Supplementary movie 2:** Live-imaging of a control-injected *Tg(mnx1:GFP)^ml2^* embryo from 20 to 38 hpf showing premature branching and truncation defects. Embryo oriented anterior to the right, dorsal to the top.

**Supplementary movie 3:** Touch response of a 48 hpf control embryo after light touch in reduced speed. Embryo uses 100 ms to swim 1 cm.

**Supplementary movie 4:** Touch response of a 48 hpf *sorcs2* KD embryo after light touch in reduced speed. Embryo uses 170 ms to swim 1 cm.

## Notes

### Competing Interest Statement

The authors have declared no competing interest.

### Summary of Updates

Correction of the first author's surname - there was a spelling mistake

